# Comprehensive Enumeration of Cancer Stem-like Cell Heterogeneity Using Deep Neural Network

**DOI:** 10.1101/2024.11.26.625418

**Authors:** Debojyoti Chowdhury, Shreyansh Priyadarshi, Sayan Biswas, Bhavesh Neekhra, Debayan Gupta, Shubhasis Haldar

## Abstract

Cancer stem cells (CSCs), a distinct subpopulation within tumors, are pivotal in driving treatment resistance and tumor recurrence, posing substantial challenges to conventional therapeutic strategies. Precise quantification and profiling of these cells are essential for improving cancer treatment outcomes. We present ACSCeND, an advanced deep neural network model accompanied by a robust workflow, specifically developed to quantify cellular compositions from bulk RNA-seq data, enabling accurate CSC profiling. By integrating bulk RNA-seq data with insights derived from single-cell RNA-seq datasets, ACSCeND effectively captures the diversity and hierarchical organization of tumor-resident cell states, alongside cell-specific gene expression profiles (GEPs). Compared to current tissue deconvolution models, ACSCeND exhibits superior performance, achieving significantly higher Concordance Correlation Coefficient (CCC) values and lower Root Mean Square Error (RMSE) across various pseudobulk and real-world bulk tissue samples. Application of ACSCeND to TCGA and PRECOG datasets reveals a strong association between CSC abundance and poorer disease-free survival outcomes, underscoring the clinical relevance of CSCs in cancer progression. Furthermore, cell-specific GEPs for distinct CSC states unveil novel molecular signatures and illuminate the origins of CSC-driven tumor heterogeneity. In summary, ACSCeND provides a powerful, scalable platform for high-throughput quantification of cellular compositions and distinct potency states within normal tissues as well as highly heterogeneous tissues, such as tumors.

## Introduction

Cancer cells possess the remarkable ability to change their phenotypes and functions, allowing them to adapt to different conditions and environments—a trait known as cancer cell plasticity^1–3^. This ability enables cancer cells to shift between different cell states, fueling tumor growth and progression. Cancer stem cells (CSCs) are believed to be key drivers of this plasticity; they can self-renew and transform into various cell types within the tumor, leading to a diverse mix of cells that develop resistance to therapies over time^4^. This same flexibility allows cancer cells to spread from the original tumor to other parts of the body (metastasis)^5^ and to relapse after a disease-free state^6,7^. Experimental evidence supports the idea that cancer cell plasticity follows a developmental hierarchy similar to that seen in normal tissue growth^8–10^. In this model, cancer stem cells sit at the top of the hierarchy, much like stem cells in healthy tissues, giving rise to various other cancer cell types through a process that mirrors normal differentiation^11,12^. Each state can come up with unique transcriptional profiles and behaviors corresponding to the tumor microenvironment (TME)^13^, thereby creating significant diversity or transcriptional heterogeneity within the tumor itself. Although the exact mechanisms that control these state changes are not fully understood, it is believed that targeting the plasticity driven by CSCs could prevent cancer from spreading or returning after treatment by eliminating the cells responsible for these adaptations.

Recent advances in large-scale and diverse scRNAseq data have provided valuable insights into the complexity of cancer, uncovering the intricate landscape of different cell states and their transitions within tumors. However, scRNAseq-based ranking methods are costly, technically challenging, and often fail to capture the full diversity of cell types present in a tumor due to selection of annotation methods, restricting their broader application^9,14^. On the other hand, bulkRNAseq-based approaches, such as mRNAsi, offer a more accessible way to measure tumor stemness but fall short in their ability to distinguish the various cell populations that make up a tumor and their interactions^15^. Trajectory inference methods might be used to arrange single cells in a developmental-trajectory like sequence that is dependent on the specific dataset^16–19^. As a result, the cells considered to be the least differentiated in one dataset might exhibit a potency level comparable to the most differentiated cells in another, complicating direct comparisons across datasets. While current deconvolution techniques^20–23^ can be used to analyze bulkRNAseq data using a signature matrix derived from different stem cell states, they will struggle to differentiate between cancer stem cells and the developmental plasticity of other tumor-associated cells. Moreover, most of these methods struggle to get a good prediction on rare and random cell fractions and therefore, might struggle in CSC enumeration. This limitation makes it harder to fully understand the dynamic interactions within the tumor environment, highlighting the need for more refined methods that can capture the full spectrum of cancer cell diversity.

To ameliorate these challenges, we developed ACSCeND (AI-based Cancer Stem Cell Profiler and Neoplasm Deconvoluter), a novel machine learning approach designed to assess cancer stem cell states along with profiling other tumor-resident cell types. This method utilizes bulkRNAseq data to effectively quantify and profile various cell states within tumors. By leveraging large-scale expression patterns and insights from scRNAseq datasets, ACSCeND accurately captures the diversity and hierarchy of cancer cell states. Our model has been rigorously validated and benchmarked using real-world data and has demonstrated remarkable accuracy in predicting qualitative CSC states. Additionally, ACSCeND has been employed to unravel the dynamics of cell-cell interactions and to investigate how CSC heterogeneity influences clinical outcomes, providing valuable insights for improving cancer research and treatment strategies.

## Results

### Overview of ACSCeND framework

The TME comprises cancer cells, stromal cells, immune cells, extracellular matrix, and vasculature, which collectively influence tumor progression^24,25^. Among these, CSCs are a rare but critical component^26,27^, uniquely capable of initiating tumor formation, driving progression, and causing relapse due to their self-renewal and differentiation capacity, and ability to adapt to environmental stresses (Figure 1A). CSCs are known to exist as a heterogeneous population, displaying diverse phenotypic and functional states influenced by genetic, epigenetic, and microenvironmental factors^28,29^. To investigate the heterogeneity within the CSC population, we classified CSCs into Pluripotent-like, Multipotent-like, and Bi/Bi/unipotent-like subpopulations based on their gene expression profiles, reflecting similarities to distinct normal stem cell states and developmental trajectories^9,15,30–33^.

**Figure 1:**
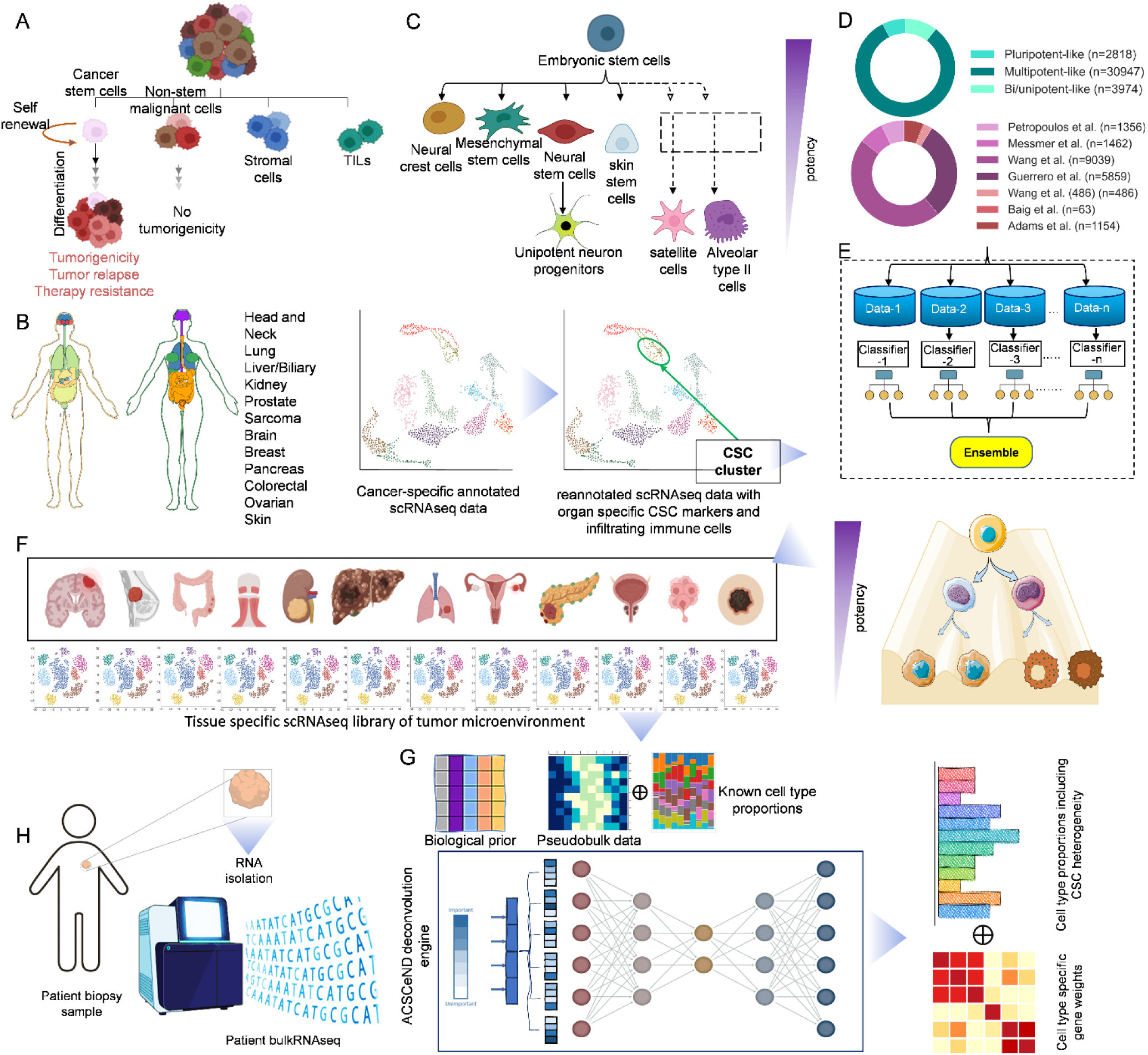
Overview of Cancer Stem Cell (CSC) Deconvolution Workflow Using ACSCEND. A. The schematic representation of developmental hierarchy model of cancer initiation, progression and evolution and the role of cancer stem cells. B. Flowchart depicting the reannotation of cancer-specific single-cell RNA sequencing (scRNAseq) data to include cancer stem cell (CSC) markers and immune cell infiltration across multiple organs. The reannotated scRNAseq data is grouped by cancer type (e.g., head and neck, lung, liver, etc.) to identify CSC clusters across tissues. Immune, stromal, and non-stem malignant cells are also highlighted for comparison. C. Schematic representation of various stem cell types taken for the training of our stem cell classifier module, including Pluripotent-like stem cells (e.g., embryonic stem cells), Multipotent-like stem cells (e.g., mesenchymal stem cells, neural stem cells, neural crest cells, and skin stem cells), and Bi/unipotent-like stem cells (e.g., Bi/unipotent-like neuron progenitors, satellite cells, alveolar type II cells). The diagram illustrates the differentiation potential of each stem cell type into distinct specialized cells or tissues, outlining their relative potencies. D. Distribution of number of cells per stem cell type and per dataset used for training the classifier. E. Representation of the ensemble model approach to classify CSC populations. Multiple datasets are generated and passed through different classifiers, which are later ensembled to identify CSC populations, grouped into Pluripotent-like-like, Multipotent-like-like, and Bi/unipotent-like-like CSC subtypes. Original architecture can be found in Extended data Figure 1. F. Diagram detailing the steps of organ-specific tumor scRNAseq library preparation to simulate pseudobulk data and known cell-type proportions. This data is then used for CSC deconvolution using the ACSCeND engine. G. Preparation of pseudobulk data, cell fraction targets and biological prior from scRNAseq library for input of ACSCeND deconvolution model. H. Schematic of the ACSCeND deconvolution engine, which deconstructs tumor bulkRNAseq data from patient biopsy samples into cell-type-specific proportions, including CSC heterogeneity. The ACSCeND model outputs include predicted proportions for various cell types (e.g., CSCs, immune cells, stromal cells) and assigns gene weights to each cell type. The ACSCeND deconvolution results display the proportions of Pluripotent-like-like, Multipotent-like-like, and Bi/unipotent-like-like CSCs across patient samples, alongside immune and non-stem malignant cells.

Determining CSC fractions from bulk RNAseq data through deep tissue deconvolution necessitates accounting for other tumor microenvironment (TME) cell types. Incorporating TME cell types ensures more accurate CSC fraction estimation while capturing the broader tumor ecosystem. To achieve this, we compiled scRNAseq data for solid tumors across 12 distinct tissues^34^. This approach reflects the tissue-dependent nature of the TME and its environmental conditions. The data were reannotated to categorize cells into immune, stromal, and malignant fractions, ensuring no overlap or contamination between cell types (Figure 1B). CSCs were specifically redefined within the malignant fraction through stemness-based clustering and marker validation (Figure 1B). We developed a classifier trained on cells representing different potency states (Figure 1C). This included a dataset of ~40 thousand cells aggregated from eight distinct sources (Figure 1D)^35–42^. Using this ensemble classifier, we categorized CSCs into Pluripotent-like-like, Multipotent-like-like, and Bi/Bi/unipotent-like-like subpopulations (Figure 1E). Integration of all these cell types resulted in a comprehensive, tissue-specific scRNAseq library of TME cell types (Figure 1F).

We developed an encoder-decoder-based deconvolution engine to estimate cell fractions within tumors from bulk RNAseq data. We designed separate engines for each solid tissue, maintaining a consistent architecture while optimizing hyperparameters specific to each tissue type. The training data for these models consisted of pseudobulk datasets generated from scRNAseq libraries, with included predefined cell fractions as targets (Figure 1G). Additionally, an scRNAseq-derived signature matrix was incorporated as a biological prior to enhance model accuracy. When applied to patient bulk RNAseq samples, our ACSCeND deconvolution engine effectively predicted CSC proportions and reconstructed detailed gene expression profiles (GEPs) (Figure 1H). The choice of deconvolution engine is determined by the tissue type from which the tumor originates.

### Prediction of CSC heterogeneity

To classify the CSCs according to different potency states, we prepared classifier modules trained with normal stem cell scRNAseq data (Extended Data Figure 1). We preferred direct validation on an external dataset to assess the model’s generalizability and determine if it was overfitting to the training data. Therefore, our analysis incorporated validation data from multiple studies (human scRNAseq data), encompassing a diverse range of cell samples (Figure 2A)^37,43–45^. Due to training sample number imbalance, we tested three approaches: training with imbalanced data, an ensemble approach and SMOTE approach (Supplementary Fig. 1). All these approaches were tested on five simple machine learning models (Extended Data Figure 2), among which the logistic regression model demonstrated superior performance, achieving near-perfect classification accuracy. The confusion matrix for the logistic regression model revealed a high accuracy in predicting all three classes (Figure 2B). Additionally, the ROC curves indicated that the model excellently managed the trade-off between sensitivity and specificity, achieving an ROC AUC of 1.000 for all potency states (Figure 2C). For the precision-recall analysis, the model attained an AUC of 1.000 for both Pluripotent-like and Bi/unipotent-like cells, and 0.992 for Multipotent-like cells, underscoring its reliability in clinical settings (Figure 2D).

**Figure 2.**
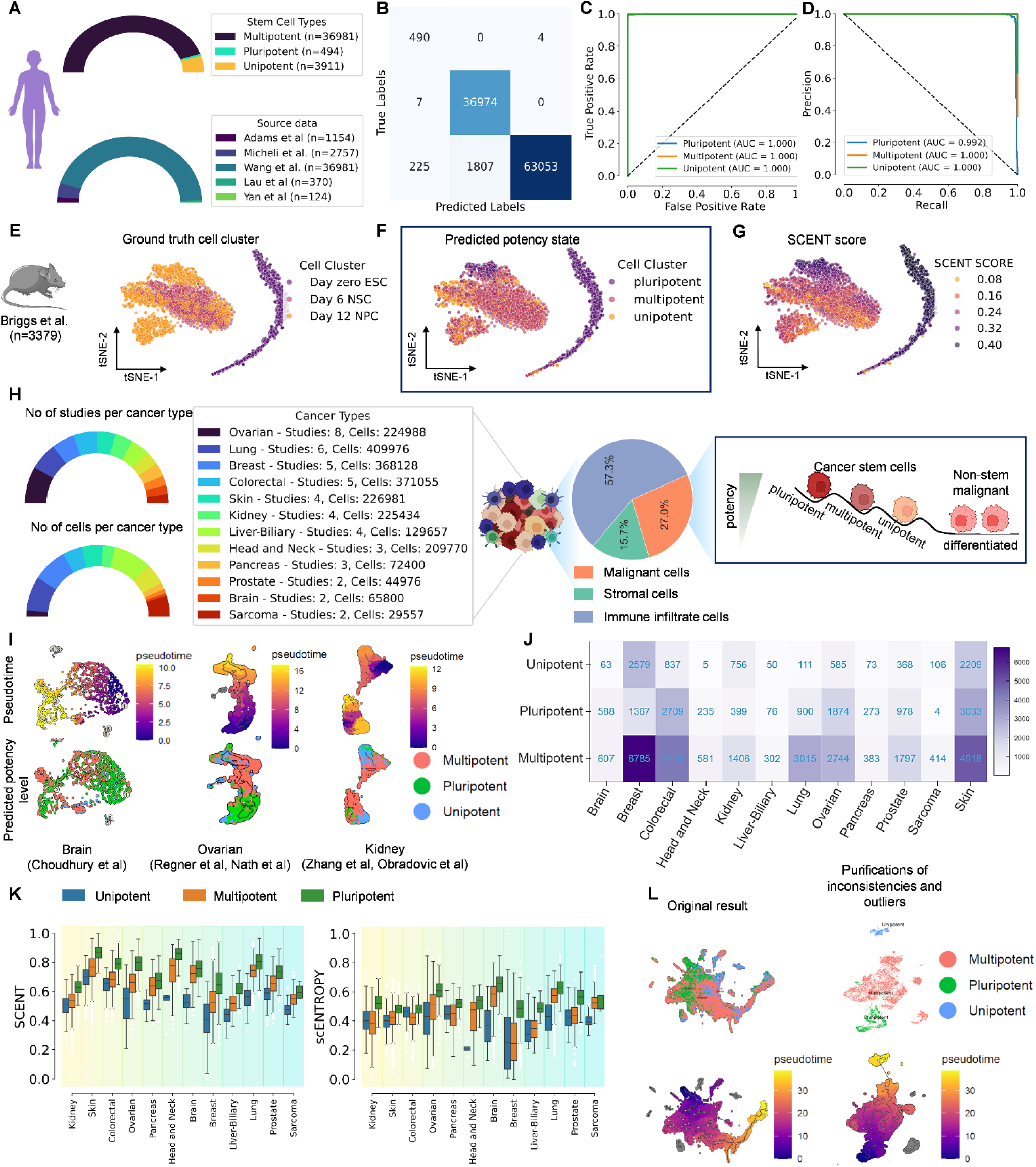
ACSCeND Classifier Module and Classification of CSC states. A. The figure represents the sources of data collected from various origins and corresponding cell types found in each data source. B. Confusion matrix between the True Labels and the Predicted Labels. C. Receiver Operating Characteristic (ROC) graph of True positive rate and False positive rate. D. Precision recall graph to illustrate the performance of the positive classes. E. Validation of ACSCeND classifier module on Mice data analysis sourced from Briggs et al. F. Prediction of ACSCEND of different potency states of stem cells on mice data. G. SCENT score calculation to quantify cellular entropy associated with cell plasticity and differentiation potential on the predicted potency states of the stem cells. Higher SCENT scores like 0.4 indicates higher potential to differentiate into multiple lineages (like pluripotency) whereas lower SCENT scores like 0.08 indicates a more specified and stable state of potency (like Bi/unipotent-like). H. Different types of cells present within the tumor microenvironment (TME). Based on studies and analysis of 12 different types of cancers, the TME is comprised of 57.3% of immune cells, 27.0% of malignant cells and 15.7% of stromal cells. The malignant cells are further comprised of different types of cancer stem cells and non-stem cancer cells. I. Pseudo-time analysis on different types of cancer, like Brain, Ovarian and Kidney, to predict the potency levels of different stem cells across time. J. The corresponding fractions of different types of stem cells are listed across different types of cancer. The x-axis represents the different cancer types while the y-axis represents the potency levels of the cancer stem cells. K. Box plots of SCENT scores and scEntropy scores of CSCs annotated by ACSCeND across 12 different types of cancers. L. The results obtained from Pseudotime analysis are further smoothened to remove the extra noise and outliers present in the result.

Validation of the model was further extended to mouse dataset, through a comparative analysis using t-SNE plots, which visually affirmed the classifier’s accuracy against ground truth labels derived from a study by Briggs et al (Figure 2E)^46^. The t-SNE plots demonstrated a clear segregation of cell types, correlating well with the predicted potency types (Figure 2F), thus confirming the model’s real-world applicability and generalizability. Further parallel comparison with application of SCENT algorithm^47^ confirmed the model’s predictive consistency across different stages of cell differentiation (Figure 2G).

As our main motive is to create scRNAseq library of TME components across 12 cancer types, we collected data from 49 studies comprising of almost 2.4 million cells (Figure 2H). These cells were pre-annotated to immune cells (57.3%), stromal cells (15.7%), and malignant cells (27%). First we annotated CSCs within the malignant cohort using a combined approach. By leveraging this reannotated CSC data, the model was able to stratify CSCs into three potency states: Pluripotent-like-like, Multipotent-like-like, and bi/Bi/unipotent-like-like CSC states. We validated the classifier model on CSC prediction through UMAP visualization and pseudotime analysis, where the model’s predictions of CSC states were confirmed across three cancer types and five datasets (Figure 2I). The UMAP plots showed clear clustering of CSCs according to their predicted potency. The alignment between the UMAP clustering and the pseudotime trajectories suggests that the model captures the temporal dynamics of CSC differentiation (Figure 2I). We applied our model on the CSC cohort to determine CSC heterogeneity in every cancer type (Figure 2J). This prediction was further confirmed by comparing our prediction results with two quantitative methods of single cell stemness calculation algorithms, SCENT^47^ and scEntropy^48^, both of which strongly agree with our predictions (Figure 2K). We further smoothened our data statistically using pseudotime, to remove inconsistencies across CSC state predictions (Figure 2L). This will ensure proper annotation of CSC states which are absolutely close to ground truth.

### Construction and Validation of ACSCeND deconvolution engine

We developed a novel encoder-decoder deconvolution model enhanced by an attention mechanism, integrating pseudobulk data with known cell type frequencies and scRNAseq signature matrices as biological priors (Extended Data Figure 3). The initial structure of the model was validated based on several parameters and hyperparameters such as attention mechanism (Supplementary Fig. 2,3), latent space dimensions (Supplementary Fig. 4), learning rate (Supplementary Fig. 5), dropout rate (Supplementary Fig. 6), batch size (Supplementary Fig. 7), and all loss functions (Supplementary Fig. 8-11, Supplementary Table 1). To assess its performance and robustness, we employed three comprehensive validation strategies. First, we partitioned pseudobulk data from four diverse datasets—PBMC 20k (Extended Data Figure 4), developmental datasets (Extended Data Figure 5)^49^, malignant datasets (Extended Data Figure 6)^50^, and mouse tissue datasets (Extended Data Figure 7)^51^— into training (50%) and testing (50%) subsets. Across all datasets, our model consistently achieved higher concordance correlation coefficients (CCC) and lower root mean square errors (RMSE) compared to state-of-the-art deconvolution methods, including TAPE^21^, SCADEN^22^, CIBERSORTx^20^, DWLS^52^, Bisque^53^, and EPIC^23^ (Extended Data Figure 4-7). These results highlight the superior predictive accuracy and reliability of our approach over existing methodologies. In the second validation strategy, we evaluated cross-dataset generalization by training the model on pseudobulk data from one source (e.g., PBMC 10k) and validating it on a distinct dataset (e.g., PBMC 20k) (Extended Data Figure 8). Similarly, training on pancreas data from Segerstople et al.^54^ and validating on pseudobulk data from Xin et al.^55^ demonstrated superior CCC and RMSE compared to benchmark methods (Extended Data Figure 8). Finally, we validated on real-world bulk RNAseq datasets with paired cell fraction data (SYD67 data,^56^ Monaco et al. data,^57^ and Newmann et al.^58^ data). Our model outperformed existing approaches in accurately estimating cell type proportions in both SYD67, and Monaco et al. data, but was at per with CIBERSORTx and TAPE on Newmann et al. data (Extended Data Figure 9). These results collectively validate the accuracy, reliability, and generalizability of our method for deconvolution tasks across diverse datasets and experimental conditions.

Accurately characterizing cellular heterogeneity in cancer remains challenging, particularly due to inconsistencies in single-cell RNA sequencing (scRNAseq) annotation methods, which vary between marker-based and reference-based approaches. These discrepancies often depend on the specific cell types being analysed, with some studies broadly identifying cell clusters while others differentiate subtypes. Notably, most scRNAseq datasets lack annotations for cancer stem cells (CSCs), further complicating analyses. To address this, we performed a uniform reannotation of tumor cell populations across datasets, ensuring consistency and accuracy (details in Supplementary Methods). UMAP analysis confirmed the validity of this approach, showing no misclassification of cell types (Supplementary Fig. 12-24). This standardized annotation framework creates tissue-specific scRNAseq libraries for tissue-specific deep deconvolution (Figure 3A).

**Figure 3.**
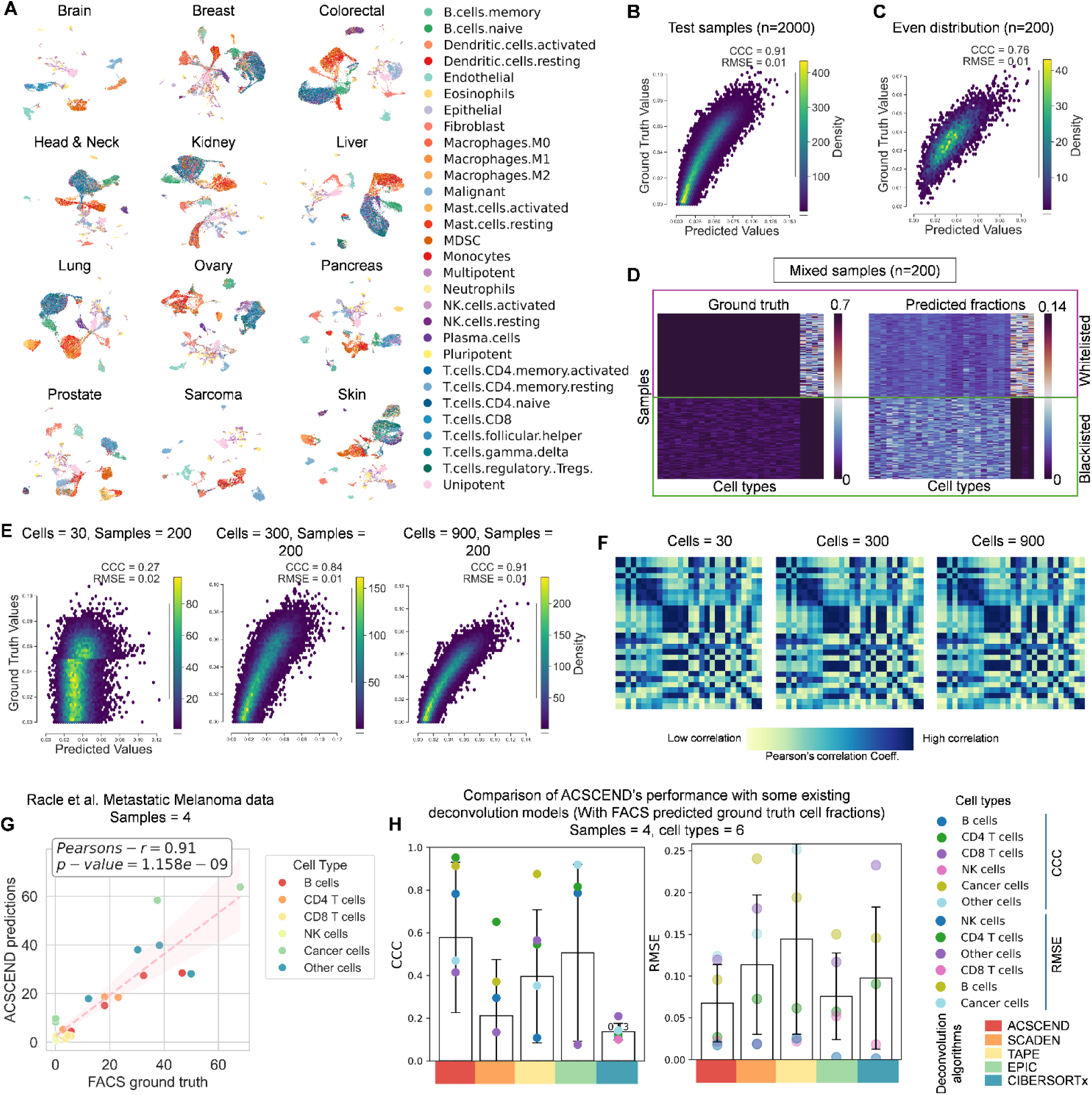
Evaluation of ACSCEND Deconvolution Model Performance Across Multiple Cancer Types and Pseudobulk Data. A. UMAP distribution of cell types across 12 different cancer datasets used for training the ACSCeND deconvolution model. The colors represent distinct cell types, as indicated in the legend. B. Performance evaluation of the ACSCeND deconvolution model on Test pseudobulk dataset with a random distribution of cell types. The CCC values are displayed within each sub-figure. C. Performance evaluation of the ACSCeND deconvolution model on pseudobulk data with an even distribution of cell types. The CCC values are displayed within each sub-figure. D. Performance evaluation of the ACSCeND deconvolution model on pseudobulk data featuring both whitelisted and blacklisted malignant and stem cell types. The model shows strong performance on pseudobulk data with blacklisted malignant and stem cell types but struggles to capture the absence of other cell types in the whitelisted group. E. Performance evaluation of the ACSCeND deconvolution model on pseudobulk data with varying cell numbers, increasing from left to right. The CCC values are displayed within each sub-figure. F. Performance evaluation of the GEPs produced by ACSCeND deconvolution model (x-axis) on pseudobulk data with varying cell numbers compared to scRNAseq ground truth (y-axis) in correlation plot. G. Performance evaluation of the ACSCEND deconvolution model on metastatic melanoma real-bulk data from Racle et al. The predicted values (Y-axis) are compared to FACS obtained ground truth cell populations (x-axis). Each cell type is shown in different color. H. Comparison of ACSCEND’s performance with some existing deconvolution models (With FACS predicted ground truth cell fractions). The bar plots represent median CCC (left plot) and RMSE (right plot). Error bars represent IQR * 1.5. Each point represents CCC or RMSE for each cell type assessed in the study.

To further examine model performance, we applied it to brain cancer data and generated pseudobulk samples with varying conditions. On 2,000 pseudobulk samples with random cell type frequencies, our model achieved a CCC of 0.91 and RMSE of 0.01 (Figure 3B), while on 200 samples with evenly distributed cell types, CCC dropped to 0.76 with an RMSE of 0.01 (Figure 3C). To assess robustness in the presence of missing cell types, we generated 200 pseudobulk samples excluding malignant cell frequencies. The model successfully identified remaining cell types, albeit with reduced performance in samples missing non-malignant cell frequencies, reflecting the higher complexity introduced by multiple missing cell types (Figure 3D). Additionally, increasing cell numbers in pseudobulk samples substantially improved model performance, with CCC rising from 0.27 (30 cells) to 0.84 (300 cells) and 0.91 (900 cells), while RMSE remained at 0.01 for larger samples (Figure 3E). Notably, gene expression profile (GEP) correlations with single-cell ground truth remained stable across varying cell numbers (Figure 3F). Validation using real-world melanoma bulk RNA-seq data from Racle et al.^59^ (n=4), which included six ground truth cell types, yielded a Pearson correlation coefficient (PCC) of 0.91 between predicted and true cell fractions (P-val = 1.158 x 10^-9^, Figure 3G). Importantly, our model demonstrated superior cell type-specific CCC (0.58 ± 0.39) and lower RMSE (0.07 ± 0.04) compared to some available deconvolution methods, indicating a significantly better performance (Figure 3H).

### Prediction of CSC dynamics at clinical settings

We applied ACSCeND on two large multi-cancer cohorts, TCGA^60^ (n = 7,020) and PRECOG^61^ (n = 6,402), to explore CSC dynamics across diverse cancer types and stages (Figure 4A). A panorama of distinct TME component cell types obtained from both datasets can be found in Figure 4B. Analysis of CSC states revealed a hierarchical pattern of Pluripotent-like > Multipotent-like > Bi/unipotent-like cells across most cancer types, consistent with the increasing plasticity and adaptability required for tumor progression (Figure 4C)^62,63^. Notably, some of the cancers such as liver cancer deviated from this trend, exhibiting a predominance of Bi/unipotent-like cells. Median CSC frequencies revealed distinct trends, with Pluripotent-like and Multipotent-like CSCs showing a multiphasic and cyclic trend across AJCC tumor stages, whereas Bi/unipotent-like CSCs demonstrated an inverse pattern of the other two states, declining in advanced stages (Figure 4D). This suggests a shift toward highly plastic CSC states as tumors progress, likely contributing to increased adaptability and resistance to therapeutic interventions. In the context of metastasis, Pluripotent-like and Multipotent-like CSCs exhibited a biphasic trend, peaking at M1A and subsequently declining at M1B and M1C stages, while Bi/unipotent-like cells showed a steady decline from M0 onwards (Figure 4E). This indicates that Pluripotent-like and Multipotent-like CSCs play critical roles during early metastatic dissemination and initial niche colonization, whereas Bi/unipotent-like CSCs may undergo depletion or differentiation in advanced metastatic sites. Tumor grade analysis also revealed a sharp increase in CSC frequencies, particularly Pluripotent-like cells, with higher grades, reflecting their involvement in driving tumor aggression and adaptability (Figure 4F).

**Figure 4.**
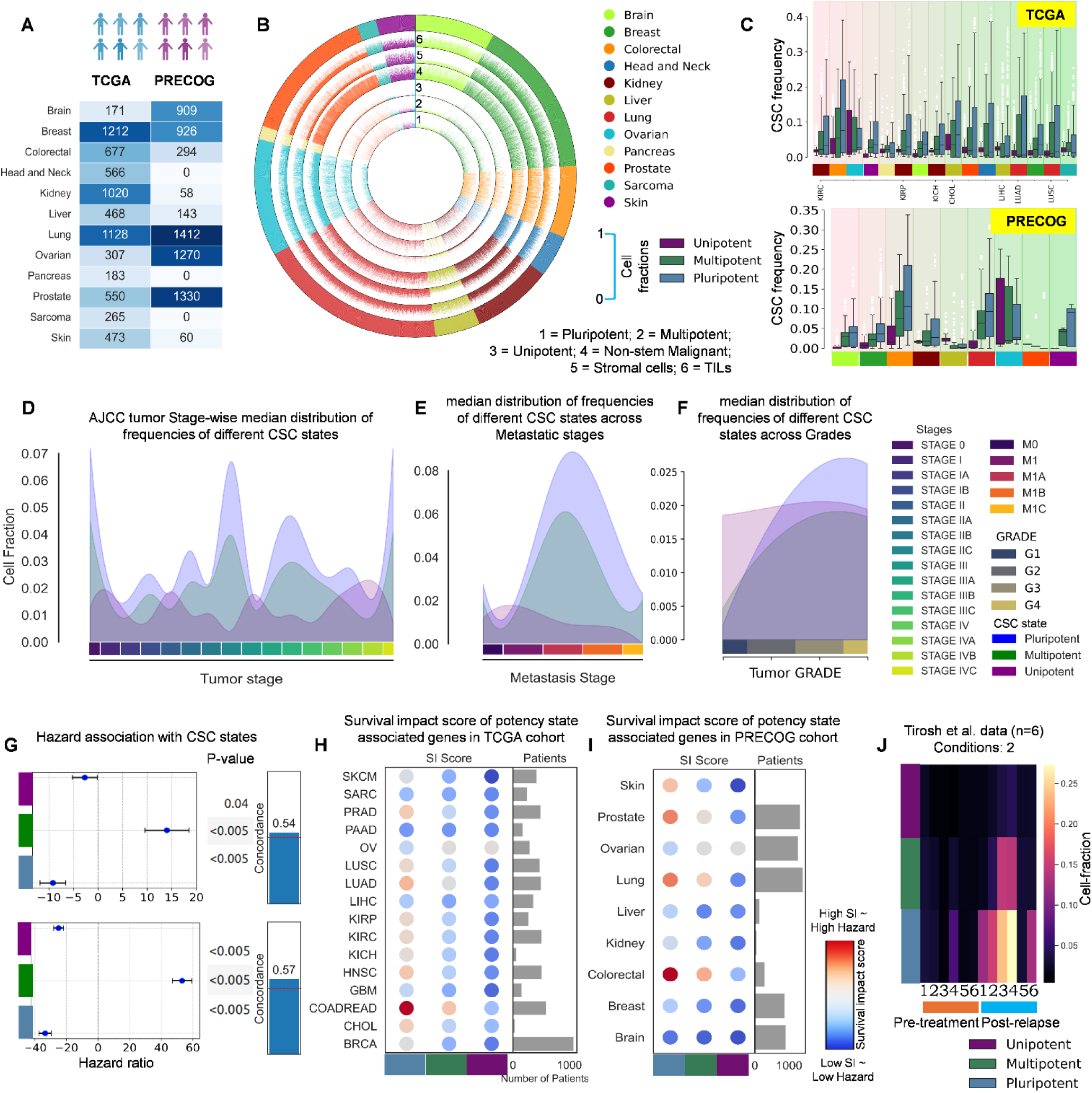
Clinical pathological associations of CSCs across multi-cancer cohorts. A. List of different quantities of cells in 12 different types of cancer for TCGA and PRECOG. B. The Circos plot illustrates the distribution of different cell types across various cancer types, with each color representing a specific cancer. Moving outward from the innermost circular segment, the plot sequentially displays the number of Pluripotent-like cells, Multipotent-like cells, Bi/unipotent-like cells, non-stem malignant cells, stromal cells, and tumor-infiltrating lymphocytes within each cancer type. C. Box plots of cancer stem cell frequencies across different types of cancers of TCGA and PRECOG listed in A. Different colors of the box plots represent different potencies of cancer stem cells. D. Plot of different types of cancer stem cell fractions at different stages of a growing tumor. Different colors of the graphs represent different types of cancer stem cells (Pluripotent-like, Multipotent-like and Bi/unipotent-like). The boxes of different colors represent different stages of cancer. A lighter colour represents a more advanced stage whereas a darker colour represents a more preliminary stage. E. Plot of different types of cancer stem cell fractions at different stages of metastasis. Different colors of the graphs represent different types of cancer stem cells (Pluripotent-like, Multipotent-like and Bi/unipotent-like). The boxes of different colors represent different metastasis stages. A lighter colour represents a more advanced stage of metastasis whereas a darker colour represents a more preliminary stage of metastasis. F. Plot of different types of cancer stem cell fractions of different tumor grades. Different colors of the graphs represent different types of cancer stem cells (Pluripotent-like, Multipotent-like and Bi/unipotent-like). The boxes of different colors represent different tumor grades. A lighter colour represents a more advanced grade of tumor whereas a darker colour represents a more preliminary grade of tumor. G. Plot of health hazards ratio associated with the different cancer stem cell states. The colored boxes along the y axis represent the different states of cancer stem cells. H. Survival impact scores of the different cancer stem cell potency state associated genes found in patients of TCGA cohort. If the colour of the score is more towards red, then it is more likely to cause more severe health hazard whereas if the colour is more towards blue then it is less hazardous. The colored boxes along the x axis represent the different states of cancer stem cells. The y axis represents the gene names. I. Survival impact scores of the different cancer stem cell potency state associated genes found in patients of PRECOG cohort. If the colour of the score is more towards red, then it is more likely to cause more severe health hazard whereas if the colour is more towards blue then it is less hazardous. The colored boxes along the x axis represent the different states of cancer stem cells. The y axis represents the cancer types. J. Figure representing fractions of cancer stem cells present before treatment and after relapse of cancer. A deeper colour represents lesser fractions of cells while a lighter colour represents the presence of more fractions of cancer stem cells present. The data was collected from the work of Tirosh et al. with 6 number of samples for both pre-treatment and post-treatment.

Survival analysis across TCGA and PRECOG cohorts in terms of hazard ratio highlighted Multipotent-like CSCs as the most prognostically adverse population, consistent to previous findings (Figure 4G)^9,14^. However, a one-time CSC prediction from initial biopsy sample only shows a static scenario whereas CSC fractions and hence their effect on survival should be dynamic. To understand those, we calculated survival impact score (SIS) for each CSC state across cancer types. This algorithm considers the contribution of top marker genes from each CSC state, and the frequency of each CSC state per sample. At this level, we find that Pluripotent-like-like CSCs are more hazardous compared to Multipotent-like-like CSCs followed by Bi/unipotent-like-like CSCs in both TCGA (Figure 4H) and PRECOG patients (Figure 4I). This contradiction is plausibly due to the ability of Pluripotent-like cells to go to a dormant or quiescent state at adverse environmental conditions.

To understand the role of CSCs in tumor relapse, we applied ACSCeND to paired pre-treatment and post-relapse bulk RNAseq data from Tirosh et al. (n=6)^64^. Post-relapse tumors showed significant enrichment of Pluripotent-like-like CSCs compared to pre-treatment samples, indicating a selective pressure for highly plastic states during therapy (Figure 4J). Interestingly, the amount of post-relapse CSCs was found to be proportional to the pre-treatment CSC load, suggesting that initial CSC burden influences the degree of CSC enrichment post-relapse. These findings highlight the dynamic and adaptive nature of CSC populations under therapeutic pressure and underscore their role in tumor recurrence, thereby providing insights into potential strategies to target CSCs for preventing relapse and improving patient outcomes.

### Reconstruction of CSC-state specific transcriptomic signatures

Our analysis of cell type-specific Gene Expression Profiles (GEPs) across tissues revealed high concordance with ground truth single-cell expression data, with most Pearson correlation coefficients (PCCs) exceeding 0.9 (Figure 5A), underscoring the reliability of our approach in capturing accurate expression patterns. Our analysis on TCGA and PRECOG data identified the top 1000 overexpressed and underexpressed genes for each stem cell type; these genes displayed distinct and robust expression profiles, reflecting genes unique to each cell type and possibly the unique mRNA markers for each cell type (Figure 5B). Further analysis of genes with continuous decreasing expression across potency states (Pluripotent-like to Multipotent-like to Bi/unipotent-like), in a pseudotemporal order. This analysis also revealed multiple experimentally validated markers consistent with their diminishing functional capacity, following the differentiation trajectories (Figure 5C). We identified the top 10 significant differentially expressed genes that might work as molecular switches to regulate transition from higher to lower potency states during tumor promoting differentiation(Figure 5D). This list includes genes involved in key pathways such as Wnt, Notch, and FOXO4, known to regulate both stem cell differentiation for lineage commitment and CSC differentiation and heterogeneity (Figure 5D).

**Figure 5.**
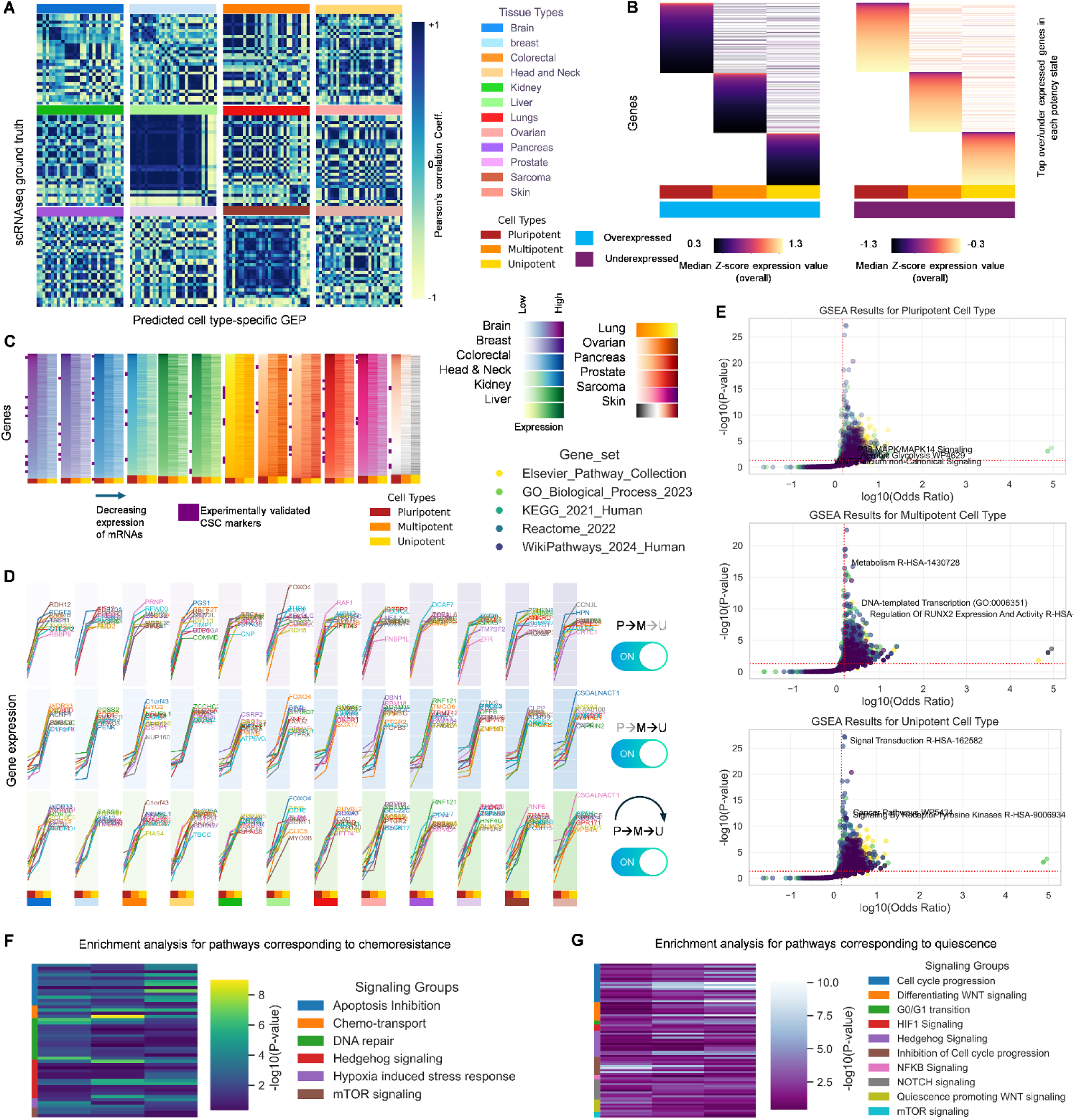
Analysis of Gene Expression Profiles (GEPs) across stem cell potency states. A. Heatmap showing Pearson correlation coefficients (PCCs) of GEPs across tissues with single-cell expression data. PCC are shown as color gradients. The color codes correspond to different tissue types. B. Heatmap depicting the top 1000 overexpressed and underexpressed genes for each stem cell type, highlighting distinct marker gene expression profiles unique to Pluripotent-like, Multipotent-like, and Bi/unipotent-like states. Expression values are shown as color gradients. The color codes correspond to different cell types. C. Heatmap with marker annotation showing the expression trajectory of genes with decreasing expression across potency states (Pluripotent-like → Multipotent-like → Bi/unipotent-like), including experimentally validated markers of differentiation. Expression values are shown as color gradients. The color codes correspond to different cell types and experimentally validated markers. D. Plots identifying the top 10 significant differentially expressed genes between potency states (Pluripotent-like vs. Multipotent-like, Multipotent-like vs. Bi/unipotent-like, Pluripotent-like vs. Bi/unipotent-like). The color codes correspond to different cell types and switch represent potency states pair E. Pathway enrichment analysis results for each potency state using Elsevier Pathway Collection, GO Biological Process 2023, KEGG 2021 Human, Reactome 2022, and WikiPathways 2024 Human databases, showing significant enrichment (odds ratio > 1.5, p < 0.05) of pathways associated with pluripotency (e.g., MAPK, Glycolysis, Wnt), multipotency (e.g., metabolism, RUNX2 regulation), and unipotency (e.g., signal transduction, RTK signaling). F. Pathway enrichment of chemoresistance-related pathways across potency states. −log10(p value) are shown as color gradients. The color codes correspond to different signalling pathways. G. Pathway enrichment of quiescence-related pathways across potency states. −log10(p value) are shown as color gradients. The color codes correspond to different signalling pathways.

We performed pathway enrichment analysis against databases such as Elsevier Pathway Collection, GO Biological Process 2023, KEGG 2021 Human, Reactome 2022, and WikiPathways Human 2024 on overall GEP set and extracted pathways important in CSC states (Figure 5E). This revealed significant enrichment (odds ratio >1.5, p < 0.05) of pathways tied to pluripotency (e.g., MAPK/MAPK14, Glycolysis, proliferative Wnt signaling), multipotency (e.g., metabolism, DNA-templated transcription, RUNX2 signaling), and unipotency (e.g., signal transduction, cancer pathways, RTK signaling), reflecting the functional stratification of potency states and their path to differentiation. Enrichment analyses of chemoresistance (Figure 5F)^65^ and quiescence-related pathways (Figure 5G)^66–68^, two important signatures of CSCs, revealed their activation across all cell types but with declining enrichment as potency decreased. Notably, DNA repair pathways were significantly enriched in Pluripotent-like-like CSCs, consistent with their enhanced genomic maintenance and chemoresistance under stress (Figure 5F). This observation aligns with the hazard ratios inferred from our analysis, where Pluripotent-like-like CSCs displayed lower hazard, likely due to increased quiescence leading to dormancy (Figure 5G). Collectively, these results demonstrate the biological relevance of GEPs in defining CSC states, identifying differentiation markers, and uncovering pathways associated with CSC differentiation, maintenance, and quiescence.

### Panorama of cancer stem cell-associated immune infiltrates

CSCs can evade immune detection and foster an anti-inflammatory, protumorigenic immune environment, aiding cancer progression^69^. For example, tumor-associated macrophages boost CSC growth through positive feedback. Our study highlights critical associations between cancer stem cell (CSC) frequencies, immune cell populations, and immunosuppressive pathways, revealing key insights into tumor heterogeneity and potential vulnerabilities for immunotherapeutic interventions.

The distribution of CSC frequencies was analyzed in the TCGA and PRECOG cohorts, stratified into hot and cold tumors based on immune infiltration status (Figure 6A). Cold tumors exhibited significantly higher CSC frequencies compared to hot tumors in both datasets, as indicated by broader distributions and elevated median frequencies. This trend was consistent across CSC states (Pluripotent-like, Multipotent-like, and Bi/unipotent-like), suggesting that cold tumors provide a niche conducive to CSC maintenance and expansion, potentially due to an immunosuppressive microenvironment.

**Figure 6.**
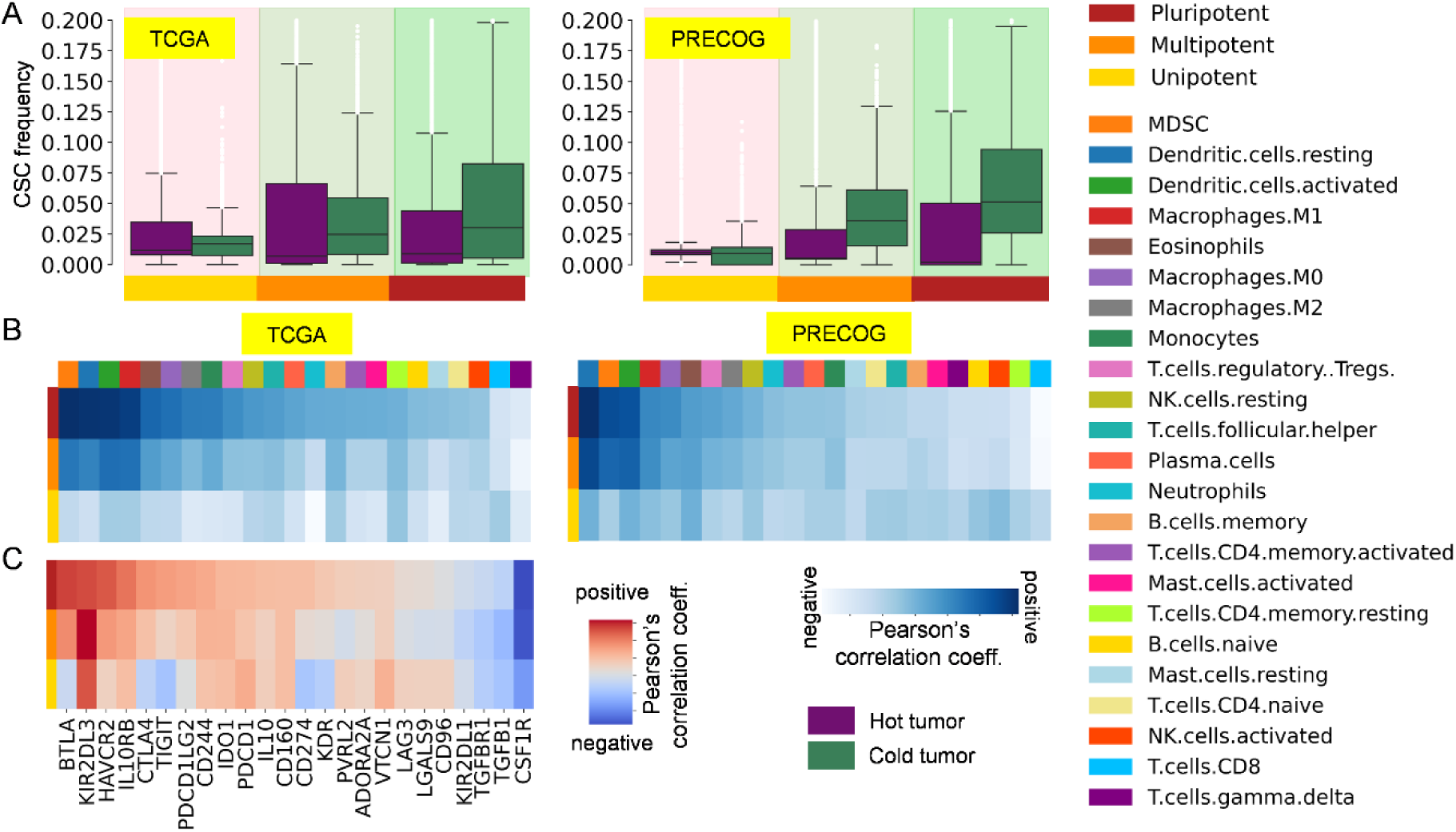
Interplay Between CSC Frequencies, Immune Cell Populations, and Immunosuppressive Pathways in Tumor Microenvironments. A. Distribution of cancer stem cell (CSC) frequencies across hot and cold tumors in the TCGA and PRECOG cohorts. Hot tumors, indicated in purple, cold tumors, shown in green, across both datasets. The x-axis groups the samples into Pluripotent-like, Multipotent-like, and Bi/unipotent-like CSC states as color coded in the legend. B. Heatmaps displaying the correlation between immune cell frequencies and CSC frequencies in the TCGA and PRECOG cohorts. Each row corresponds to a specific immune cell population, while columns represent individual tumor samples. Blue color intensity reflects a stronger positive correlation, whereas lighter blue indicates weaker associations or no correlation. C. Correlation between the expression of immunosuppressive genes and CSC frequencies across tumors. Red regions signify a positive Pearson correlation coefficient while, blue regions denote a negative correlation.

Correlation analysis of immune cell frequencies and CSC frequencies revealed distinct relationships across both TCGA and PRECOG cohorts (Figure 6B). Immune populations such as MDSCs, M1/M2 macrophages, and resting dendritic cells showed positive correlations with CSC frequencies, whereas cytotoxic immune populations, including CD8+ T cells and natural killer (NK) cells, exhibited weaker or negative correlations. These findings suggest that CSC-rich tumors are characterized by an immune-suppressive landscape, potentially driven by Tregs and tolerogenic dendritic cells, which may shield CSCs from immune-mediated elimination.

Further investigation into the relationship between immunosuppressive gene expression and CSC frequencies uncovered significant correlations (Figure 6C). Genes encoding immune checkpoint molecules, including PDCD1 (PD-1), HAVCR2 (TIM-3), and BTLA, were positively associated with CSC frequencies, emphasizing the role of immune evasion pathways in promoting CSC persistence. Interestingly, cytokine and immune-modulating genes such as TGFB1 and CSF1R also showed positive correlations, further implicating these pathways in the regulation of CSC states. Conversely, certain genes with negative correlations may represent potential regulators that suppress CSC expansion.

These findings have critical implications for immunotherapeutic strategies targeting CSCs. The association of CSCs with immunosuppressive immune cell populations and checkpoint gene expression highlights potential avenues for disrupting CSC-immune interactions. Immune checkpoint inhibitors (e.g., anti-PD-1, anti-TIM-3) may not only reinvigorate cytotoxic immune responses but also disrupt the protective niches that foster CSC survival. Consistent with available litteratures our analysis also suggests that therapies targeting TGF-β or macrophage-associated pathways (e.g., CSF1R inhibitors)^70^, or immunotherapies with γδ-Tcells^71^ could attenuate the immunosuppressive microenvironment, reducing CSC frequencies and their ability to evade immune surveillance.

## Discussions

Our AI-driven framework, ACSCeND, represents a groundbreaking advancement in cancer stem cell profiling and tumor deconvolution. By leveraging bulk RNA-sequencing (bulkRNAseq) data alongside insights from single-cell RNA-sequencing (scRNAseq) datasets, ACSCeND provides a comprehensive approach to dissecting the intricate cellular composition of tumors. Its innovative dual encoder-decoder architecture integrates latent cell-type representations with an attention mechanism, enabling precise quantification of CSC states and other tumor-resident cell types. The tissue-specific model design, which dynamically selects optimal configurations based on dataset complexity, ensures robust performance across diverse tumor microenvironments. ACSCeND’s strength lies in its advanced deconvolution engine, which significantly outperforms existing methods. This stems from its flexible architecture, which adapts seamlessly to datasets of varying complexities, distinguishing it from models that are narrowly optimized for specific contexts. Compared to other bulkRNAseq deconvolution models, ACSCeND demonstrates superior performance in capturing the dynamic spectrum of CSC states and other cellular populations within tumors. Across multiple datasets, including pseudobulk and real bulk data, ACSCeND consistently achieves higher CCC and lower RMSE, underscoring its reliability and robustness. Moreover, ACSCeND outperforms other models in time-complexity analysis as well, as it takes lower time for data processing and prediction in total, compared to other models (Extended Data Figure 10).

Beyond cancer profiling, ACSCeND holds transformative potential for broader applications in biomedical research. Its ability to deconvolute complex cellular mixtures with high precision makes it a valuable tool for studying the tumor microenvironment, immune cell dynamics, and the cellular responses to therapeutic interventions. Moreover, the model’s adaptability allows its extension to developmental biology, where understanding cell-state transitions and differentiation processes is critical. ACSCeND’s validated performance across human and mouse datasets further supports its utility for cross-species comparisons, offering new insights into conserved and species-specific cellular mechanisms. These features position ACSCeND as a versatile tool with far-reaching implications beyond oncology, including regenerative medicine and developmental biology. Furthermore, application of ACSCeND is feasible for both raw count and normalized data (TPM, CPM or FPKM), independent of gene expression quantification methods (RNAseq or microarray) making it a broadly applicable deconvolution algorithm.

Despite its many strengths, ACSCeND is not without limitations. As with any supervised machine learning approach, the quality and diversity of the training data heavily influence its performance. While ACSCeND effectively captures a broad range of cell states, underrepresented developmental and functional states may constrain its accuracy. Expanding the training dataset to include greater diversity and underrepresented cell types could address these gaps and enhance the model’s generalizability. Additionally, the model occasionally struggles to recognize the absence of certain cell types, particularly when these are preselected for analysis. Addressing this limitation will require further refinement of the model’s handling of dataset-specific nuances. Moreover, the reliability of ACSCeND’s predictions depends on the quality of input data, emphasizing the importance of continuous validation against experimental benchmarks.

In conclusion, ACSCeND marks a significant step forward in the field of cancer biology, offering a powerful and adaptable framework for profiling CSC dynamics and tumor heterogeneity. By bridging the methodological gap between bulkRNAseq and scRNAseq, ACSCeND provides a scalable, accurate, and versatile tool for understanding complex cellular ecosystems. As the training framework expands and the model undergoes further refinements, ACSCeND is poised to revolutionize our understanding of tumor biology, CSC behavior, and their implications for therapeutic strategies, ultimately paving the way for more effective cancer treatments and precision medicine approaches.

## Methods

### Single-Cell RNA Sequencing Data Acquisition and Quality Control

The single-cell RNA sequencing (scRNA-seq) gene expression matrix for various cancer types was sourced from the publicly available Weizmann Institute’s 3CA dataset (Provide Supplemetary Table). Initial preprocessing and quality control were conducted using the Seurat package (v5.1.0) in R. Cells expressing fewer than 200 genes or more than 6,000 genes, as well as cells with a mitochondrial gene fraction exceeding 10%, were excluded from downstream analyses to remove low-quality cells. Suspected doublets were identified and filtered using the DoubletFinder package (v2.0) in R^72^. The optimal pK parameter was determined by iterating through potential values using the paramSweep function, followed by summarization and selection of the maximum BCmetric from the output of the find.pK function. Homotypic doublets were accounted for by adjusting the expected doublet rate (nExp_poi.adj) based on the proportion of homotypic cells. Doublets were then removed before proceeding to further analysis.

### Data Integration and Dimensionality Reduction

After the initial quality control, the gene expression matrix was normalized using the Seurat package (v5.1.0) in R^73^. Normalization was performed with the NormalizeData function, utilizing the LogNormalize method and a scale factor of 10,000 to standardize gene expression levels across all cells. To identify genes with the highest variability across cells, the FindVariableFeatures function was applied using the variance-stabilizing transformation (vst) method, and the top 2,000 highly variable genes were selected. The dataset was subsequently scaled using the ScaleData function, which standardizes the expression of each gene across all cells. Principal Component Analysis (PCA) was then conducted using the RunPCA function on the identified variable genes to get the 20 PCs. To address potential batch effects arising from technical variations across different experimental conditions, batch correction was performed using the RunHarmony function of the Harmony package. FindNeighbors function is then utilized to construct the nearest neighbor graph and subsequent clustering of cells into distinct groups was performed using the Louvain algorithm implemented in Seurat’s FindClusters function, with a resolution parameter of 0.3.

### Cell Type Annotation and Cluster Identification

Cell type annotations for malignant cells, fibroblasts, epithelial cells, and endothelial cells were preserved as provided in the original dataset. For the annotation of immune cell populations, the SCINA package^74^ was utilized, leveraging marker genes specified in the LM22 matrix by Newman et al^75^. Myeloid-derived suppressor cell (MDSC) marker genes were selected from Alshetaiwi et al.^76^ The annotation of cancer stem cells (CSCs) involved a multi-step manual approach (See supplementary Methods). First, malignant cells were isolated based on the original annotations and subjected to clustering using the Louvain algorithm, as implemented in Seurat. Clusters were then visualized using violin plots to assess the expression levels of curated cancer stem cell markers specific to each cancer type. For each cluster, the expression patterns of these markers were compared to identify those with elevated levels, indicative of a potential stem cell phenotype (Supplementary Table 2).

### The classifier module

Experimentally annotated datasets were aggregated from multiple publicly available sources for training our classifier model. To ensure the consistency and quality of the data, we calculated the SCENT score for all samples. The SCENT score was utilized to smoothen the dataset by removing any samples whose original annotations were not in alignment with the score. This process ensured that the data used for model training reflected accurate biological classifications. Following this refinement, a class imbalance was observed in the data, where Multipotent-like samples constituted approximately eight times the number of Pluripotent-like and Bi/unipotent-like samples. To address this, rather than discarding the data, we employed an ensemble learning strategy. Specifically, eight distinct ensemble models were trained. These models shared common Pluripotent-like and Bi/unipotent-like samples, but each incorporated a unique subset of Multipotent-like samples, thereby balancing the data across the different models. Before model training, the entire dataset was converted to rank space. This approach assigned relative ranks to gene expression levels, thereby removing batch effects, mitigating the influence of extreme values and outliers, and reducing the risk of overfitting. The rank space conversion also normalized gene expression differences between samples, facilitating robust downstream analysis. Subsequently, the rank-transformed data was log2-transformed and standardized to z-scores.

To further ensure robustness and prevent overfitting, we did not employ traditional k-fold cross-validation, which can introduce bias when biological replicates are present in both the training and testing sets, leading to the issue of double-dipping. Instead, our models were tested on completely independent datasets collected from entirely different sources. This strategy allowed for an unbiased evaluation of model performance, ensuring that the trained models could generalize effectively to novel data. We didn’t perform SCENT-based smoothening on the test data prior to prediction, maintaining the integrity of the unseen test data for accurate evaluation. We developed five different models, and for subsequent analysis, we selected the one that demonstrated the highest accuracy on the independent test dataset. We implemented the following machine learning classifiers, along with their parameters for model training:

1. Logistic Regression - The parameters used are : penalty=‘l2’, *, dual=False, tol=0.0001, C=1.0, fit_intercept=True, intercept_scaling=1, class_weight=None, random_state=None, solver=’lbfgs’, max_iter=100, verbose=0, warm_start=False, n_jobs=None, l1_ratio=None.
2. Decision Tree Classifier - The parameters used are : criterion=‘gini’, splitter=‘best’, max_depth=None, min_samples_split=2, min_samples_leaf=1, min_weight_fraction_leaf=0.0, max_features=None, random_state=None, max_leaf_nodes=None, min_impurity_decrease=0.0, class_weight=None, ccp_alpha=0.0, monotonic_cst=None.
3. RandomForestClassifier - The parameters used are : _estimators=10, *, criterion=‘gini’, max_depth=None, min_samples_split=2, min_samples_leaf=1, min_weight_fraction_leaf=0.0, max_features=‘sqrt’, max_leaf_nodes=None, min_impurity_decrease=0.0, bootstrap=True, oob_score=False, n_jobs=None, random_state=None, verbose=0, warm_start=False, class_weight=None, ccp_alpha=0.0, max_samples=None, monotonic_cst=None.
4. MLPClassifier - The parameters used are : hidden_layer_sizes=[512, 256], activation=‘relu’, *, solver=“sgd”, alpha=0.0001, batch_size=‘256’, learning_rate=‘constant’, learning_rate_init=0.001, power_t=0.5, max_iter=100, shuffle=True, random_state=None, tol=0.0001, verbose=False, warm_start=False, momentum=0.9, nesterovs_momentum=True, early_stopping=False, validation_fraction=0.1, beta_1=0.9, beta_2=0.999, epsilon=1e-08, n_iter_no_change=10, max_fun=15000
5. XGBClassifier - The parameters used are : n_estimators=2, max_depth=20, learning_rate=1, num_class=3, objective=‘multi:softmax’

### Cancer Stem Cell Classification and Validation

The annotated cancer stem cells (CSCs) were classified into three distinct subtypes using a previously established classification model. To validate the robustness of this classification, we computed the Single-cell Energy Conversion Efficiency of Neural Transcription (SCENT) score for each annotated sample. Box plot visualizations (refer that box plot) demonstrated the distribution of SCENT scores across the identified subtypes, further supporting the classification results.To gain insights into the developmental trajectories of these CSCs, pseudotime analysis was conducted using the Monocle3 package. This analysis models the progression of cells along a pseudo-temporal axis, reflecting their differentiation state. The resulting pseudotime values were segmented into three distinct clusters via K-means clustering, capturing the major developmental pathways of CSCs. Samples exhibiting concordance between their assigned stem cell subtype and pseudotime cluster were prioritized, ensuring consistency and robustness for downstream deconvolution analyses.Following the comprehensive annotation of CSCs, stromal cells, and immune cells, a balanced dataset for deconvolution was prepared. For each cell type, a minimum of 1,000 cells was retained where available, to ensure statistical robustness. To mitigate batch effects across the final dataset, the pycombat_seq function from the inmoose package was employed, achieving uniformity across samples and facilitating accurate deconvolution analysis.

### The deconvolution engine

#### Data Preparation

The datasets used in this study include raw count data for single-cell RNA sequencing (scRNAseq), pseudobulk RNA sequencing, and real bulk RNA sequencing. The scRNAseq data provides the expression of individual cells, annotated with corresponding cell types in a separate file. The pseudobulk RNA-seq data is generated by aggregating scRNAseq profiles from multiple cells, simulating bulk RNA-seq profiles. Additionally, ground truth cell fractions corresponding to each pseudobulk profile are provided, representing the true proportion of each cell type in the simulated bulk samples. The cell types were mapped using a cell annotation file. Each dataset consists of raw counts for gene expression, with rows representing genes and columns representing either individual cells (for scRNAseq) or pseudobulk samples. The final dataset used for training and validation includes the pseudobulk expression profiles as features and the ground truth cell fractions as labels.

#### Input Data dimension preprocessing

The initial step in our model is the preprocessing of pseudobulk data denoted as *X* ∈ *R^H^ ^X^ ^N^* where H is the number of genes and N is representing the number of samples. Specifically, we transform the values by standard procedure of log normalization to mitigates the effect of outliers and allows for a more Gaussian-like distribution of the values. After log transformation, we standardize the data using standard scalar to produce z-score normalization.

The Gene Expression Profiles G ∈ *R^H^ ^X^ ^P^* must be aligned with the pseudobulk dataset to ensure that both the datasets are compatible for downstream analysis. P represents the number of cell types. The transpose of the standardized pseudobulk, along with the targets which are the corresponding cell fraction datasets are divided into training, validation and test sets to facilitate model training and evaluation. The data are split into 60% training, 20% validation, and 20% test datasets. The train, validation and test datasets and the signature matrix are then converted into pytorch tensors.

#### Model Architecture

The model is an Encoder-Decoder that aims to deconvolute bulk RNA sequencing (RNAseq) data to infer cell-type proportions (cell fractions) and estimate gene expression profiles (GEPs) specific to each cell type.

The model takes the Pseudobulk gene expression dataset as input *X* ∈ *R^N^ ^X^ ^H^*, where N is the number of samples in the training dataset and H is the number of genes present in each sample (For simplicity, the transpose of the fraction of pseudobulk used as training data is represented by X). Each row represents the expression levels of the genes in a particular biological sample. The task of deconvolution is to predict the proportion of different cell types in each sample, represented by the target matrix *C* ∈ *R^N^ ^X^ ^P^* corresponding to the training dataset, where P is the number of cell types. The relationship between gene expression and cell fractions is governed by a Gene Expression Profiles G, which defines how strongly each gene is expressed in each cell type.

The deconvolution goal is summarized as:

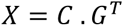

This equation shows that pseudobulk matrix X is approximately equal to the product of the cell fractions Y and the transposed Gene Expression Profiles *G^T^*. In this context, G captures the unique gene expression profiles of each cell type.

1. Encoder The encoder is designed to reduce the dimensionality of the input training data X while learning complex gene relationships that contribute to predicting cell fractions. This is achieved by passing X through multiple fully connected layers and an attention mechanism. The training data is first transformed through fully connected layers that progressively reduce the dimensionality. Each layer extracts meaningful patterns and interactions between genes that are relevant for predicting cell types. These layers capture non-linear dependencies between genes, which are important in capturing complex biological relationships. A multiheaded self-attention mechanism (heads = 8) is applied to the second layer of the encoder to enhance the model’s ability to focus on the most relevant gene interactions. For each head attention scores are calculated using:

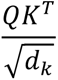

And hence the output of the attention layer is calculated by:

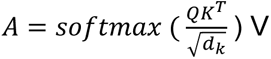 Where *Q*, *K*, *V* ∈ *R*^2048^ are the query, key, and value vectors derived from the input and *d_k_* is a scaling factor. This self-attention step allows the model to weigh the contributions of different genes dynamically. In biological terms, some genes are more critical than others in distinguishing cell types, and attention helps the model identify these key genes. Mathematically, attention computes the weighted interactions between all genes, where the weights are learned based on the model’s focus on specific gene-gene interactions. The attention output is passed through additional fully connected layers with ReLu activation functions and dropouts to form the latent representation.
2. Latent Representation After passing through the encoder, the gene expression data is transformed into a latent representation. This latent space plays a central role in the model’s ability to deconvolute gene expression data and predict cell fractions. The latent space can be thought of as a compressed, biologically meaningful representation of the input data. It captures the essential features of the gene expression profiles that are most relevant for distinguishing between different cell types. In biological systems, while the expression of thousands of genes contributes to the cell’s function, only a subset of these genes may be critical for defining a particular cell type. The latent space helps the model focus on this key subset. This compression reflects how the complexity of gene expression patterns can be reduced to a few key patterns that correspond to different cell types. The transformation into latent space can be described as:

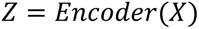 Where *Z* ∈ *R^N^ ^X^ ^d^* is the latent space representation, and d is the latent space dimensionality.
3. Decoder The decoder takes over to predict the cell fractions and reconstruct the training pseudobulk data. The latent representation Z is transformed through several fully connected layers to predict the cell fractions *C_pred_* ∈ *R^N^ ^X^ ^P^*,which represent the proportions of different cell types in each sample. To ensure the predicted cell fractions are valid (i.e., non-negative and summing to 1 for each sample), a SoftMax function is applied to the output of the final fully connected layer:

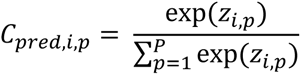

After predicting the cell fractions, the model also reconstructs the pseudobulk matrix by multiplying the predicted cell fractions *C_pred_* with the known Gene Expression Profiles G. This step ensures that the predicted cell fractions are biologically plausible by checking if they can accurately reconstruct the original training data.

The Gene Expression Profiles (GEPs) are learnt through the decoder using gradient descent method:

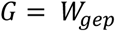

Where *W_gep_* is a trainable parameter representing gene expression signatures.

#### Loss function

The model’s learning is guided by a combined loss function which is the combination of four different loss functions:

1. The primary loss term is the mean squared error (MSE) between the predicted cell-fractions *C_pred_* and the true cell fractions *C_true_*.
2. The pseudobulk reconstruction loss is to ensure that the predicted cell fractions and signature matrix can accurately reconstruct the pseudobulk gene expression.
3. From the predicted GEPs, the loss between reconstructed pseudobulk profiles and true profiles has also been calculated. The ability to accurately reconstruct pseudobulk data allows us to validate the model’s predictions against actual experimental data, ensuring that the inferred cell-type compositions align with biological realities. It also provides a mechanism to assess the model’s performance in generating realistic gene expression profiles, which is vital for downstream analyses, such as differential expression studies or pathway analyses that rely on accurate expression data.
4. KL Divergence loss measures how one probability distribution diverges from a second expected probability distribution. It is often used as a regularization term to encourage the predicted distribution to be close to a prior distribution:

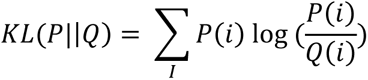 Where P is the predicted distribution and Q is the prior distribution.

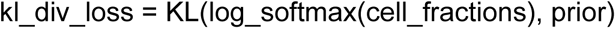

This loss encourages the model to make the predicted cell fractions adhere to a specified prior distribution.

#### Model performance validation matrics

Initially the model’s performance was checked on the validation split and further the accuracy was checked on the test dataset to look for any overfitting. After that the accuracy was checked on a completely independent dataset to confirm the applicability of the model. We used two commonly used matrices for this purpose: CCC (Lin’s concordance correlation coefficient) and RMSE (Root mean square error). The Concordance Correlation Coefficient (CCC) is used to measure the agreement between predicted and true proportions. The CCC ranges from −1 to 1, with 1 indicating perfect concordance. A higher CCC value and a lower RMSE suggest a better deconvolution performance. These are defined as follows:

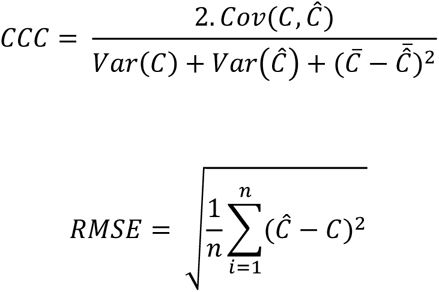

Where:

- C is the set of true values (e.g., actual cell fractions).
- *Ĉ* is the set of predicted values (e.g., predicted cell fractions).
- *Cov*(*C*, *Ĉ*) is the covariance between C and *Ĉ*.
- Var(C) and Var(*Ĉ*) are the variances of C and *Ĉ* respectively.

*C̄* and 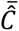 are the means of C and *Ĉ* respectively.

### Performance Evaluation

To rigorously evaluate the robustness and predictive accuracy of our deconvolution model, we employed a comprehensive validation framework comprising three distinct strategies: pseudo-bulk data validation, cross-dataset generalization, and real-world bulk RNA-seq dataset testing. In the first approach, pseudo-bulk datasets were generated from four diverse sources, including PBMC 20k, developmental datasets, malignant datasets, and mouse tissue datasets. Each dataset was partitioned into training and testing subsets in a 50:50 ratio to ensure balanced evaluation. This strategy assessed the model’s capacity to accurately infer cell-type proportions under controlled conditions. The second approach tested cross-dataset generalization by training the model on pseudo-bulk data derived from one dataset and validating its performance on a distinct dataset. For instance, pseudo-bulk data from PBMC 10k were used for training, while PBMC 20k served as the validation dataset. Similarly, pancreas data from Segerstople et al. were used for training, and pseudo-bulk data from Xin et al. were employed for validation. To assess performance in real-world scenarios, the third approach applied the model to bulk RNA-seq datasets with paired cell fraction annotations, including SYD67, Monaco et al. and Newman et al. These datasets provided experimentally determined ground-truth cell-type proportions, enabling precise evaluation of the model’s predictions. Across all datasets, performance metrics such as the Concordance Correlation Coefficient (CCC) and Root Mean Square Error (RMSE) were computed, and the results were benchmarked against state-of-the-art deconvolution tools such as TAPE, SCADEN, CIBERSORTx, DWLS, Bisque, and EPIC.

To further investigate robustness, we applied the model to brain cancer pseudo-bulk datasets generated under diverse conditions. These included 2000 samples with randomized cell-type frequencies and 200 samples with uniformly distributed frequencies. Additionally, we evaluated model performance in scenarios with missing cell types by generating 200 samples excluding malignant cell fractions and another 200 samples excluding non-malignant cell fractions. The impact of varying cell numbers in pseudo-bulk samples was systematically examined by increasing constituent cell numbers from 30 to 300 and then to 900 cells. This variation enabled us to assess the model’s sensitivity to sample composition. Performance in these scenarios was further analyzed by correlating gene expression profiles (GEPs) with median single-cell expression values across different cell counts. Finally, the model was applied to melanoma bulk RNAseq data from Racle et al., which included six annotated cell types with established ground truth. Model predictions were compared using metrics such as the Pearson Correlation Coefficient (PCC), CCC, and RMSE with performance evaluated against leading deconvolution methods. This comprehensive evaluation framework, incorporating both synthetic and real-world datasets, established the accuracy, generalizability, and robustness of our model across diverse biological contexts.

For deconvolution performance on pseudo-bulk and real bulk datasets with known ground truth, we benchmarked TAPE, Scaden, CIBERSORTx, DWLS, EPIC, and Bisque. The details of the benchmarking procedures are explained below, and the hyperparameter tuning information can be found in Supplementary Informations.

### Clinical Data and Analysis

We retrieved expression and survival data (n = 6,402) from the PRECOG database across nine cancer types. and obtained gene expression data (n = 7,020) from TCGA^77^ for cancers including KIRC, COADREAD, OV, SKCM, PAAD, KIRP, GBM, KICH, CHOL, PRAD, HNSC, LIHC, LUAD, BRCA, LUSC, and SARC through the Broad GDAC repository. These cancer types were selected because they were used to train the deconvolution model. Corresponding clinical data for TCGA cancers were sourced from cBioPortal^78^ to facilitate clinical analysis.

We performed an in-depth evaluation of Pluripotent-like, Multipotent-like, and Bi/unipotent-like stem cell frequencies across various clinical and pathological categories within the TCGA dataset. These included AJCC pathologic tumor stage (n = 4,879), pathological M stage (PATH_M_STAGE; n = 4,707), and tumor grade (GRADE; n = 1,841). This combined analysis of TCGA cancers provides insights into stem cell frequency variations across critical clinical and pathological parameters.

### Survival analysis and survival impact score

A Cox Proportional Hazards analysis was performed using lifeline package to assess the association of Pluripotent-like, Multipotent-like, and Bi/unipotent-like cell frequencies with overall survival (OS). This analysis aimed to estimate the extent to which the hierarchical stem cell states influenced patient risk.

For survival association calculation, the hazard values for individual genes were derived by leveraging the Cox Proportional Hazards Model, as implemented in the lifelines Python package. Specifically, the CoxPHFitter function was utilized to model the relationship between gene expression levels and overall survival outcomes, incorporating both survival time and censoring status for each patient. For each gene, a separate univariate Cox regression analysis was performed to estimate its hazard ratio, quantifying the extent to which the gene’s expression level influenced patient risk. A hazard ratio greater than 1 indicates an increased risk of adverse survival outcomes with higher expression of the gene, while a ratio less than 1 suggests a protective effect. These hazard ratios were subsequently log-transformed to enable downstream aggregation and analysis, facilitating the integration of individual gene-level effects into a coherent framework for evaluating cumulative survival risks. It allowed us to systematically assign hazard scores to genes, which were then used as inputs for quantifying the collective impact of gene expression patterns on patient survival outcomes within specific cell types.

Building on the gene-specific hazard values, we developed a Survival Impact Score (SIS) to quantify the contribution of cell-type-specific hazards to patient survival. To compute SIS, the proportional abundances of cell types in each patient were first normalized, ensuring the fractions accurately reflected relative cell-type compositions. For a given cell type, its SIS was calculated as the product of its normalized fraction and the aggregated hazard values of its associated genes, where the aggregated hazard was determined by summing the pre-assigned hazard values across all genes expressed within the cell type. This methodology resulted in a patient-cell-type matrix of SIS values, effectively capturing the cumulative risk contributions of specific cell types. To further understand the relationship between these cell-type-specific contributions and overall survival, Pearson correlation analysis was employed. This involved calculating correlation coefficients to assess the strength and direction of association between the SIS for each cell type and patient survival times. Statistical significance was evaluated for each correlation, with p-values log-transformed to enhance interpretability and identify meaningful associations. It enabled the identification of cell types that were significantly associated with survival outcomes, linking the cumulative risk contributions of cell-type-specific gene expression to overall patient risk profiles in a mechanistically insightful manner.

### Analysis of Immune-CSC interactions

We analyzed immune cell frequencies computed by a deconvolution model across the TCGA and PRECOG datasets, focusing on their relationship with stem cell frequencies. For each cancer type, we calculated the Pearson Correlation Coefficient (PCC) between the frequencies of Multipotent-like, Pluripotent-like, and Bi/unipotent-like stem cells and 33 immune cell types. To investigate the relationship between stem cell frequency and immunoinhibitory gene expression, we utilized genes associated with immunoinhibition identified by Lu et al. The PCC was computed between the expression levels of these genes and stem cell frequencies across the merged TCGA and PRECOG datasets, providing insights into stem cell-mediated immune modulation.

To assess the impact of immunotherapy on cancer stem cell frequencies, we analyzed datasets from metastatic melanoma patients treated with immune checkpoint inhibitors, including those from Hugo et al.^79^, and Van Allen et al.^80^. Stem cell frequencies were compared before and after treatment, enabling us to evaluate potential therapeutic effects on cancer stem cell dynamics in response to immunotherapy.

### Statistics and reproducibility

Determining the appropriate sample size for the deconvolution analysis is challenging, as no established methods currently exist. Therefore, we selected the sample size based on previously published datasets to demonstrate the effectiveness and performance of our method. No data were omitted from the analysis. Since no experimental data were collected, replication and randomization are not applicable. Statistical methods used for hypothesis testing are described in each figure legend. Group allocations were based on information from previously published datasets and were not modified by us. To replicate the findings, please refer to the Source Data file provided.

Downstream statistical analyses were conducted using common Python packages, specifically versions 3.11.6 or 3.12.0. and R packages for version R 4.3.1. The figures were generated using Python packages: matplotlib, seaborn, and pandas. Pseudobulk data were generated using R-package SimBu. Additional packages like anndata and tqdm were used for developing our method.

## Data Availability

This study includes scRNAseq datasets from different sources. All these datasets, their source article, and other details with links to the original data can be found at https://github.com/SML-CompBio/ACSCEND. The TCGA data can be obtained from https://www.genome.gov/Funded-Programs-Projects/Cancer-Genome-Atlas, https://gdac.broadinstitute.org/, or https://www.cbioportal.org/. PRECOG cancer data can be obtained from https://precog.stanford.edu/.

## Code Availability

R and Python packages for running ACSCeND with the pre-trained model are available at https://github.com/SML-CompBio/ACSCEND. ACSCeND is freely available for non-profit academic use at https://github.com/SML-CompBio/ACSCEND.

## Acknowledgments

We thank S.N. Bose National Centre for Basic Sciences and Ashoka University for support and funding. S.H. thanks DST SERB Core Research Grant for funding. D.C. thanks DBT for funding and support through the DBT-JRF fellowship. B. N and D. G are thankful to the Mphasis F1 Foundation for their support. The funders had no role in the study design, data collection and analysis, decision to publish or preparation of the manuscript.

## Author Contributions

S.H. and D.G. conceived the study. D. C, S. P, S. H, and D. G designed the study. D.C, S. P, and S. B curated the data. S. B and D. C developed the deconvolution model. S.P developed the classifier model. S. P and D. C have done the reannotation of cancer scRNAseq library. D. C, and S. P performed the data analysis. D. C, S. P, B. N and D. G wrote the manuscript. S. P and B. N built the packages. All the authors took part in interpreting the findings and reviewing the manuscript.

## Correspondence

Correspondence should be addressed to Debayan Gupta and Shubhasis Haldar. Requests for codes should be addressed to Shreyansh Priyadarshi and Bhavesh Neekhra.

