## Supplementary Information for "Comprehensive Enumeration of Cancer Stem-like Cell Heterogeneity Using Deep Neural Network"

#### Supplementary Figure Contents

### Supplementary Figures

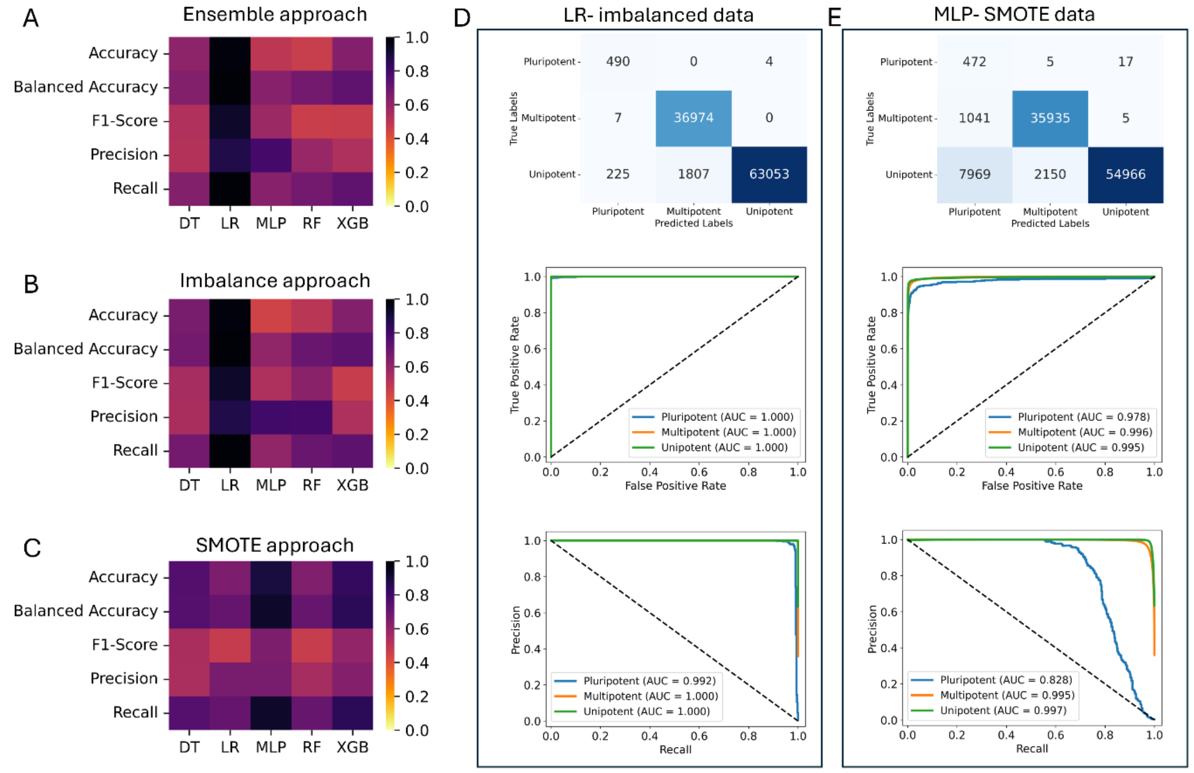

Supplementary Figure 1 Evaluation of Machine Learning Models on Imbalanced and SMOTE technique of data handling for stem cell classification.

(A) Heatmap comparing the performance of five machine learning models—Decision Tree (DT), Logistic Regression (LR), Multi-Layer Perceptron (MLP), Random Forest (RF), and XGBoost (XGB)—on imbalanced data using an ensemble approach. Performance metrics evaluated include Accuracy, Balanced Accuracy, F1-Score, Precision, and Recall, with a color scale from 0 (lowest performance) to 1 (highest performance). Logistic Regression (LR) exhibits the highest values across multiple metrics, indicating strong performance in the ensemble approach. (B) Heatmap comparing the same models (DT, LR, MLP, RF, XGB) evaluated on imbalanced data without any resampling methods. Performance metrics (Accuracy, Balanced Accuracy, F1-Score, Precision, Recall) are visualized in a similar color scale. Overall, performance appears lower compared to the ensemble approach, with balanced accuracy and recall particularly affected by the class imbalance. (C) Heatmap showing the comparison of model performance on data balanced with SMOTE (Synthetic Minority Over-sampling Technique). The five models (DT, LR, MLP, RF, XGB) are evaluated with the same set of metrics. Although performance improves for certain models with SMOTE, the results suggest that some metrics, such as balanced accuracy and recall, remain below optimal levels, particularly for models other than Logistic Regression. (D) Confusion matrix, ROC-AUC (Receiver Operating Characteristic - Area Under Curve), and Precision-Recall curve for the Logistic Regression model evaluated on imbalanced data. The confusion matrix indicates high specificity and sensitivity, particularly for the "Multipotent" and "Unipotent" classes, with almost perfect ROC-AUC scores (1.000) across all classes. The Precision-Recall curve also shows high precision and recall values, confirming strong classification performance despite class imbalance. (E) Confusion matrix, ROC-AUC, and Precision-Recall curve for the Multi-Layer Perceptron model evaluated on SMOTE-balanced data. The confusion matrix reveals improved performance compared to the imbalance approach, although misclassifications are observed, particularly in the "Pluripotent" class. The ROC-AUC scores are slightly lower than those of the Logistic Regression model, with values of 0.978, 0.996, and 0.995 for the respective classes. The Precision-Recall curve shows high recall for all classes but slightly lower precision for the "Pluripotent" class.

The Logistic Regression model using the ensemble approach was chosen as the final model for stem cell classification due to its consistently high performance across all metrics in both the ensemble and imbalance approaches, as well as its ability to handle class imbalances effectively. The ensemble approach demonstrated superior balanced accuracy, F1-score, and recall, essential for reliable classification across the diverse cell types. This selection ensures a robust and generalizable model for accurately classifying stem cell types in imbalanced datasets.

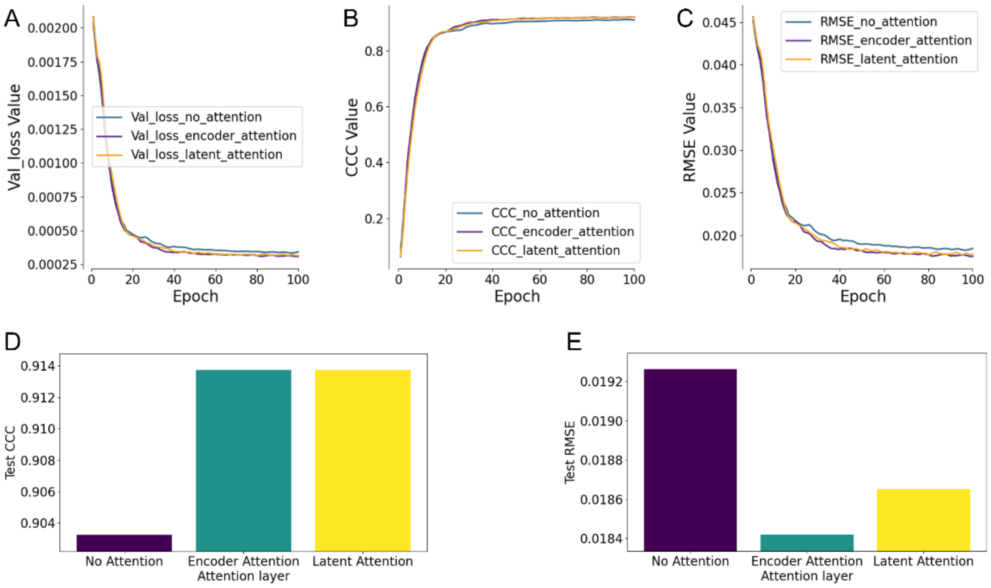

Supplementary Figure 2 Optimization of attention position.

(A) validation loss across training epochs for three scenarios: without attention, with encoder attention, and with latent attention. All three configurations show a decreasing trend in loss over epochs, but the encoder and latent attention configurations exhibit slightly lower loss values towards the end of the training. (B) progression of the CCC values for each attention mechanism across training epochs. Encoder and latent attention configurations show a similar and slightly better improvement in CCC over epochs, compared to the model without attention, indicating their positive impact on model performance. (C) reduction in RMSE across epochs for each attention setting. Encoder and latent attention configurations lead to slightly lower RMSE values than the model without attention, indicating improved accuracy in predictions with attention mechanisms. (D) CCC values on the test dataset for each attention mechanism. Both encoder and latent attention layers demonstrate significantly higher CCC compared to no attention, with a marginal difference between encoder and latent attention configurations. (E) RMSE values on the test dataset for each attention mechanism. Encoder attention achieves the lowest RMSE, followed by latent attention, with the model without attention having the highest RMSE, indicating a clear performance boost with attention layers.

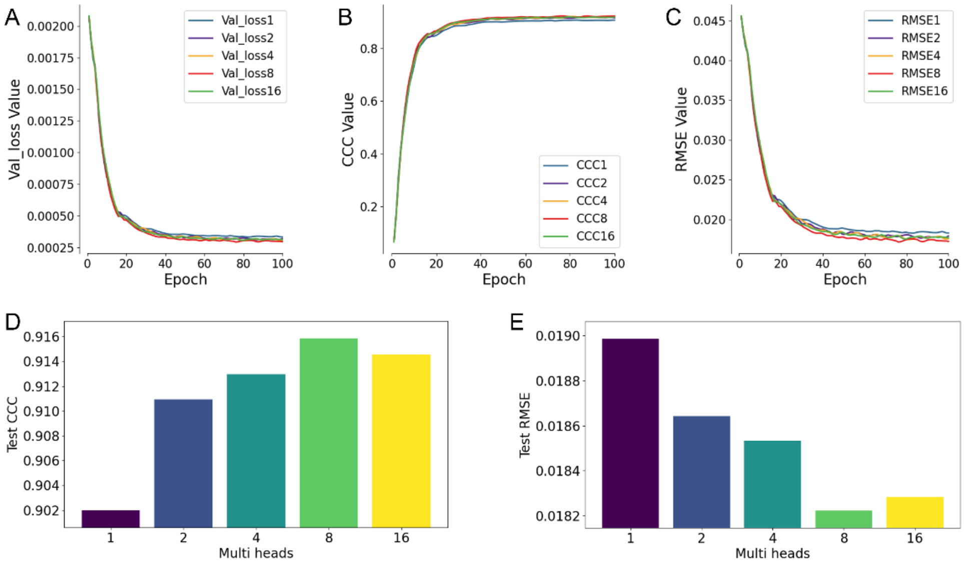

Supplementary Figure 3 Attention Head Optimization for ACSCEND Deconvolution Model

(A) the validation loss over training epochs for different numbers of attention heads (1, 2, 4, 8, and 16). As the number of heads increases, the model's validation loss decreases steadily across epochs, converging to a lower final loss for all configurations. This indicates that increasing the number of heads helps capture more diverse patterns in the data, leading to improved model training. (B) Concordance Correlation Coefficient (CCC) values over training epochs for different numbers of attention heads. Higher CCC values indicate better agreement between predicted and true cell-type proportions. The graph suggests that as the training progresses, attention configurations WITH 8 AND 16 reach similar levels of CCC, demonstrating the model's HIGHER effectiveness in accurately predicting cell proportions. (C) validation RMSE values over training epochs for different numbers of attention heads. The decreasing RMSE values over time signify the model's improving prediction accuracy as it learns from the training data. All attention configurations follow a similar trend, reaching comparable RMSE levels by the final epochs, indicating that increasing the number of heads provides marginal gains in reducing error beyond a certain point. (D) CCC values on the test dataset for each attention HEAD. Both encoder and latent attention layers demonstrate significantly higher CCC compared to no attention, with a marginal difference between encoder and latent attention configurations. (E) RMSE values on the test dataset for each attention mechanism. Encoder attention achieves the lowest RMSE, followed by latent attention, with the model without attention having the highest RMSE, indicating a clear performance boost with attention layers.

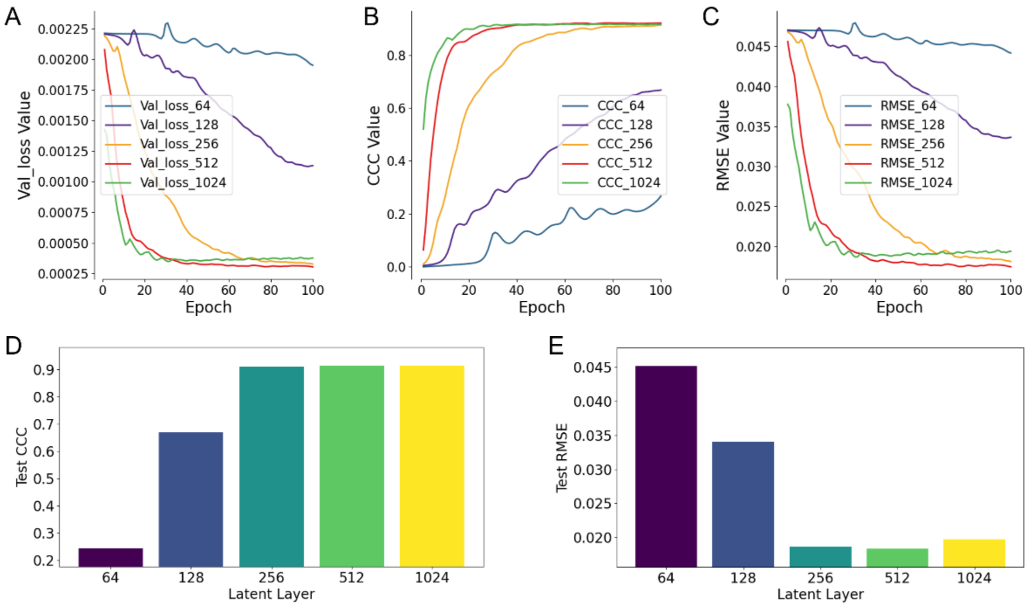

Supplementary Figure 4 Optimization of latent space dimensions.
(A) The validation loss curves demonstrate that as the latent layer size increases, the model achieves lower and more stable loss values. Notably, with 512 latent units, the validation loss converges to a minimal level quickly and stabilizes, indicating a good balance between model complexity and generalization. Beyond 512 units, the loss does not improve significantly, suggesting that larger dimensions offer diminishing returns. (B) This panel shows a clear improvement in CCC as the latent layer size increases, with 512 units achieving near-optimal correlation between predictions and actual values. The CCC curve for 512 units reaches a high plateau early in the training, indicating that this dimension size allows the model to capture the necessary information effectively without the added complexity of larger sizes. (C) The RMSE curves reveal that the model with 512 latent units consistently achieves one of the lowest error levels across all epochs. This demonstrates that this latent size is sufficient to minimize prediction errors while maintaining training stability, making it a strong choice for model accuracy. **(D)** In the test CCC comparison, the model with 512 latent units achieves a value comparable to the larger sizes (1024), indicating that it effectively captures correlations without overfitting or requiring additional complexity. This supports the choice of 512 units as an optimal balance between performance and model size. (E) The test RMSE comparison highlights that the model with 512 units achieves one of the lowest RMSE values, similar to larger sizes like 1024. This further reinforces that a 512-dimensional latent space offers minimal errors on unseen data while avoiding unnecessary computational overhead.

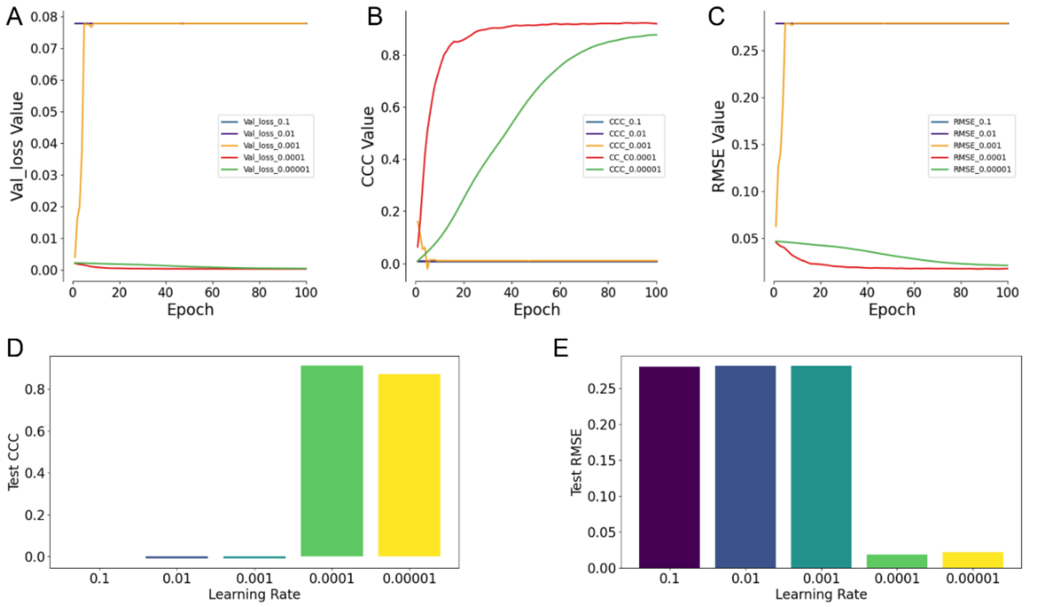

Supplementary Figure 5 Learning Rate Optimization for ACSCEND Deconvolution Model.

(A) validation loss over training epochs for different learning rates (0.1, 0.01, 0.001, 0.0001, and 0.00001). As the graph indicates, higher learning rates (0.1 and 0.01) result in significantly high and stagnant loss values, showing no improvement over epochs. On the other hand, smaller learning rates (0.001, 0.0001, and 0.00001) demonstrate better convergence and reach lower loss values quickly, indicating effective learning by the model. (B) CCC values over training epochs for each learning rate. Learning rates of 0.001 and 0.0001 achieve higher CCC values earlier in the training process and maintain this performance over epochs. In contrast, higher learning rates (0.1 and 0.01) show no progress in CCC values, while a very low learning rate (0.00001) displays a slower rise in CCC but eventually stabilizes at a reasonable level. (C) RMSE values over training epochs for each learning rate. Higher learning rates (0.1 and 0.01) exhibit consistent, elevated RMSE values, indicating that these rates hinder model performance. Lower learning rates (0.0001 and 0.00001) successfully reduce RMSE over epochs and converge to minimal error values, highlighting their effectiveness in improving model accuracy. The smallest learning rate (0.00001) also reduces RMSE but at a slower pace. (D) CCC values on the test dataset for each learning rate mentioned in the figure legend. Both 0.0001 and 0.00001 learning rate demonstrate significantly higher CCC compared to the higher learning rates (≥0.001). (E) RMSE values on the test dataset for each learning rate. A learning rate 0.0001 achieves the lowest RMSE, followed by 0.00001, with the model with 0.1, 0.01, and 0.001 having higher RMSE, indicating a clear performance boost with 0.0001 learning rate.

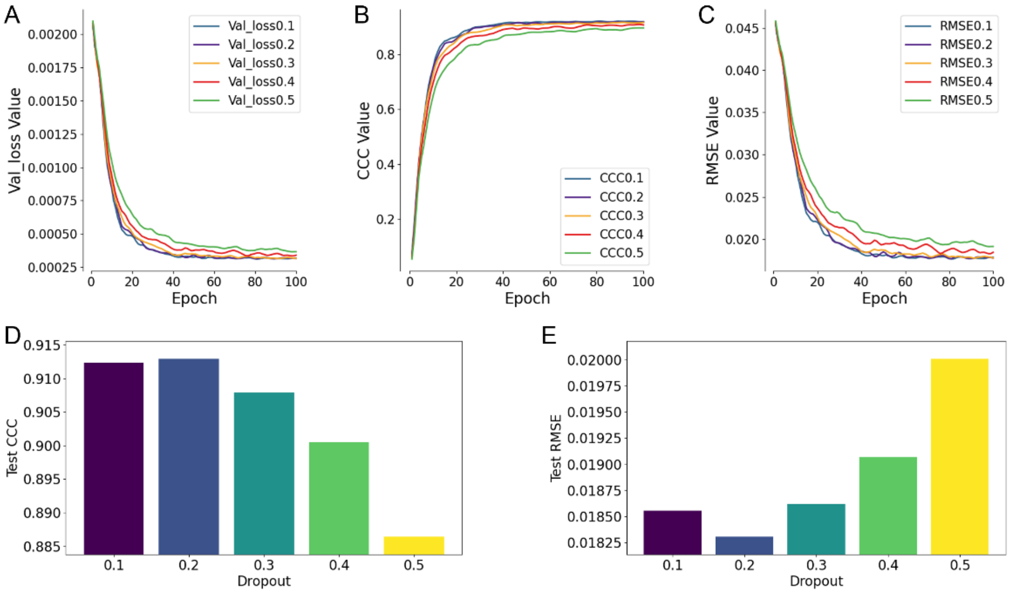

Supplementary Figure 6 Dropout rate optimization for ACSCEND Deconvolution model.

(A) the effect of different dropout rates (ranging from 0.1 to 0.5) on validation loss over 100 epochs. It is evident that a lower dropout rate (0.1 or 0.2) leads to consistently lower validation loss throughout training. As dropout rates increase beyond 0.3, the loss remains higher, indicating that larger dropouts may hinder the model's ability to learn effectively. (B) The CCC curves indicate that lower dropout rates (0.1 and 0.2) result in higher CCC values and faster convergence. As the dropout rate increases to 0.4 or 0.5, the CCC values stabilize at a lower level, suggesting a decline in predictive performance at these higher rates. (C) RMSE values across epochs for each dropout rate. Lower dropout rates (0.1 and 0.2) yield the lowest RMSE values, indicating reduced errors in the model's predictions. Higher dropout rates lead to increased RMSE, suggesting poorer performance. (D) test CCC bar chart reveals that the model achieves the highest test CCC with a dropout rate of 0.1 and 0.2. As the dropout rate increases to 0.4 and 0.5, the CCC decreases, implying that higher dropout values negatively impact the model's correlation with actual values. (E) test RMSE for each dropout rate. The lowest RMSE is observed at a dropout rate of 0.1, with the error increasing as the dropout rate rises. This confirms that a lower dropout rate minimizes prediction errors on the test set.

The results indicate that a dropout rate of 0.1 or 0.2 achieves the best trade-off between validation loss, CCC, and RMSE. Dropout rates higher than 0.2 lead to a decline in model performance, making 0.1 or 0.2 the optimal choice for the ACSCEND deconvolution model.

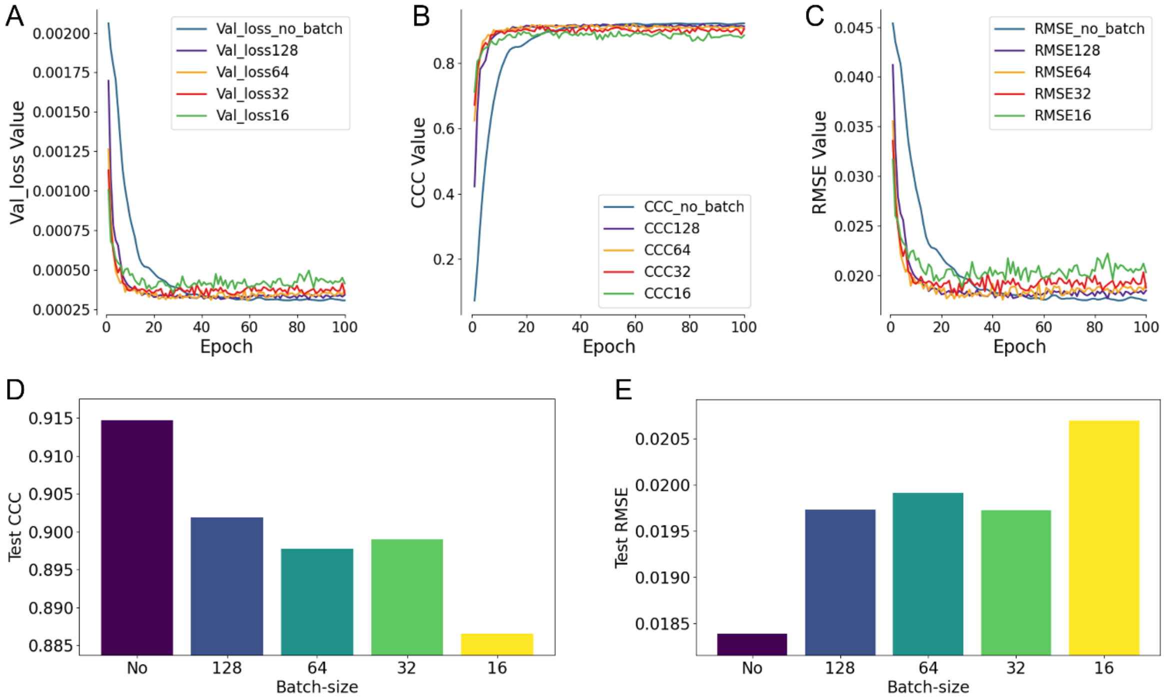

Supplementary Figure 7 Batch Size Optimization in Ascend Deconvolution Model

(A) Validation loss values over 100 epochs for different batch sizes (No_batch, 128, 64, 32, and 16). The model without batch processing (No_batch) shows a rapid initial decline in loss, achieving the lowest overall validation loss by the end of training. In contrast, smaller batch sizes (16 and 32) exhibit greater fluctuations, indicating instability during training. (B) Concordance correlation coefficient (CCC) values on the validation set over 100 epochs for each batch size configuration. The No_batch setup achieves consistently higher CCC values earlier in training and maintains the best performance, suggesting a stronger correlation between predicted and actual values. Models trained with batch sizes (especially 128) converge slower and plateau at lower CCC values, highlighting a reduction in predictive accuracy as batch size increases. (C) Root mean square error (RMSE) values on the validation set across epochs for various batch sizes. The No_batch configuration consistently maintains the lowest RMSE, while larger batch sizes (e.g., 128) exhibit initially high RMSE values that converge slowly, underscoring the stability and precision of the model without batching. (D) Test set CCC for each batch size, showing that the No_batch configuration reaches the highest CCC value (~0.915), outperforming all batch configurations. Smaller batch sizes (32 and 16) yield slightly lower CCC values, while larger batch sizes (128 and 64) show a significant decrease in test CCC, indicating poorer alignment between predicted and true outputs. (E) Test set RMSE for each batch size, demonstrating that the No_batch model achieves the lowest RMSE (~0.0185), with error progressively increasing as batch size decreases, reaching a peak for the batch size of 16.

This trend suggests that although smaller batches may benefit model generalization in certain cases, the No_batch setup provides superior error minimization. These results collectively highlight that omitting batch processing (No_batch) yields the most favorable balance of low validation loss, high CCC, and low RMSE, reflecting its stability and accuracy in the deconvolution model's performance on both validation and test data.

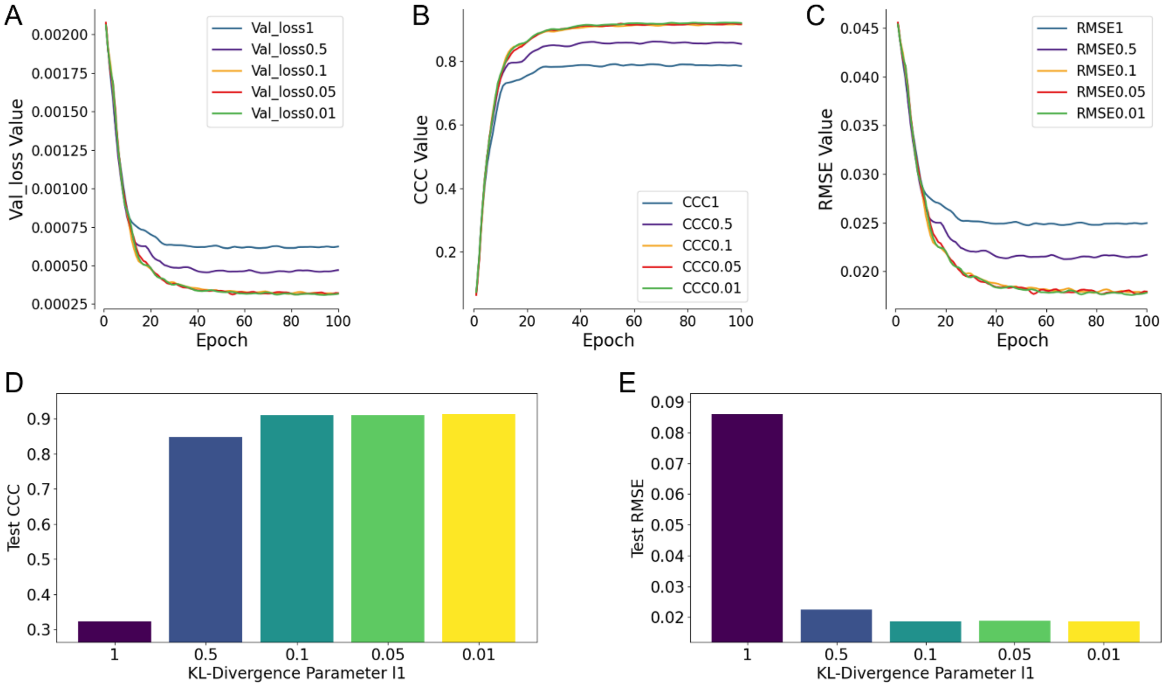

Supplementary Figure 8 KL Divergence Loss Coefficient Optimization in Ascend Deconvolution Model
(A) Validation loss values across 100 epochs for different KL divergence loss coefficients (λ): 1, 0.5, 0.1, 0.05, and 0.01. Lower λ values (especially 0.01 and 0.05) achieve the lowest validation loss, indicating more stable convergence and better fit to the data, while larger λ values (e.g., 1) result in higher loss throughout training. (B) Concordance correlation coefficient (CCC) values on the validation set over epochs for different λ values. Lower values of λ (0.1, 0.05, and 0.01) yield the highest CCC, maintaining strong predictive accuracy, while higher values (1 and 0.5) result in lower CCC, demonstrating weaker agreement between predicted and true values. (C) Root means square error (RMSE) values on the validation set across epochs for each λ. The smallest λ values (0.01 and 0.05) achieve the lowest RMSE, indicating that a minimal KL divergence constraint (or none) allows the model to minimize error effectively, whereas higher λ (e.g., 1) leads to significantly higher RMSE. (D) Test set CCC for each λ, showing that the smallest λ values (0.01, 0.05, and 0.1) produce the highest CCC values, with minimal differences between them. Conversely, larger values (1) drastically reduce CCC, suggesting poorer predictive performance on the test set. (E) Test set RMSE for each λ, demonstrating that λ=0.01 achieves the lowest RMSE, followed closely by λ=0.05 and 0.1. Larger values (1) lead to substantial increases in RMSE, emphasizing that minimal KL divergence constraint improves the model's accuracy. These results collectively highlight that a KL divergence coefficient of 0.01 provides the best balance of low validation loss, high CCC, and low RMSE, supporting the selection of λ=0.01 for optimal performance in the deconvolution model.

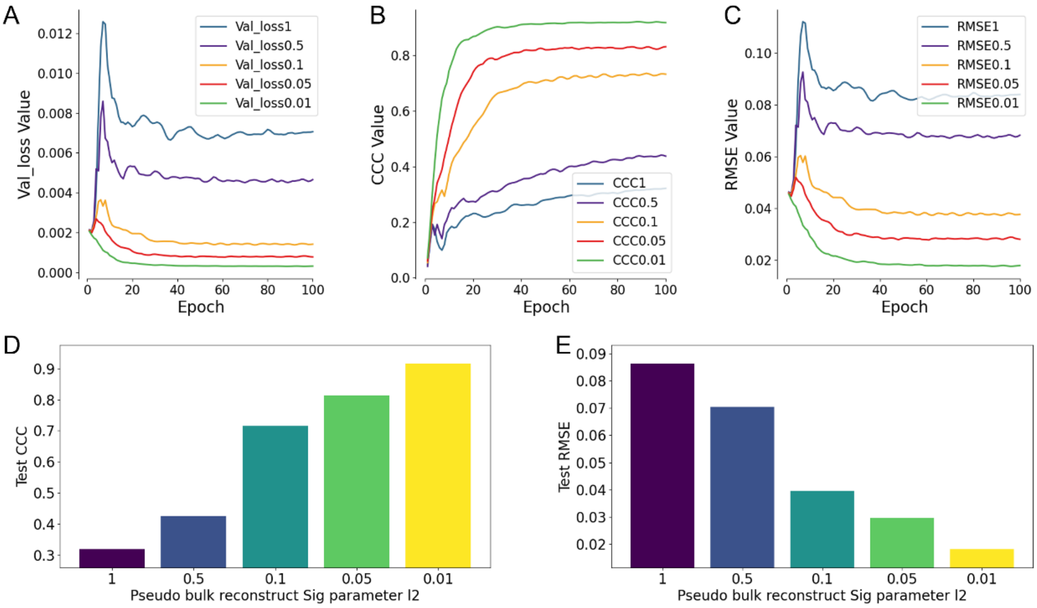

Supplementary Figure 9 Pseudo-bulk Reconstruction Loss (using Signature Matrix) Coefficient Optimization in Ascend Deconvolution Model (l2)

(A) Validation loss values across 100 epochs for different Pseudo-bulk reconstruction loss (using Signature Matrix) coefficients (λ): 1, 0.5, 0.1, 0.05, and 0.01. Lower λ values (especially 0.01) achieve the lowest validation loss, indicating more stable convergence and better fit to the data, while larger λ values (e.g., 1) result in higher loss throughout training. (B) Concordance correlation coefficient (CCC) values on the validation set over epochs for different λ values. Lower values of λ (especially 0.01) yield the highest CCC, maintaining strong predictive accuracy, while as values of λ are getting higher, it results in lowering of CCC values, demonstrating weaker agreement between predicted and true values. (C) Root means square error (RMSE) values on the validation set across epochs for each λ. The smallest λ values (0.01) achieve the lowest RMSE, indicating that a Pseudo-bulk reconstruction loss (using Signature Matrix) constraint (or none) allows the model to minimize error effectively, whereas higher λ (e.g., 1) leads to significantly higher RMSE. (D) Test set CCC for each λ, showing that the smallest λ values (0.01) produce the highest CCC values. Conversely, larger values (1) drastically reduce CCC, suggesting poorer predictive performance on the test set. (E) Test set RMSE for each λ, demonstrating that λ=0.01 achieves the lowest RMSE, followed by λ=0.05 and 0.1. Larger values (1) lead to substantial increases in RMSE, emphasizing that minimal Pseudo-bulk reconstruction loss (using Signature Matrix) constraint improves the model's accuracy. These results collectively highlight that a Pseudo-bulk reconstruction loss (using Signature Matrix) coefficient of 0.01 provides the best balance of low validation loss, high CCC, and low RMSE, supporting the selection of λ=0.01 for optimal performance in the deconvolution model.

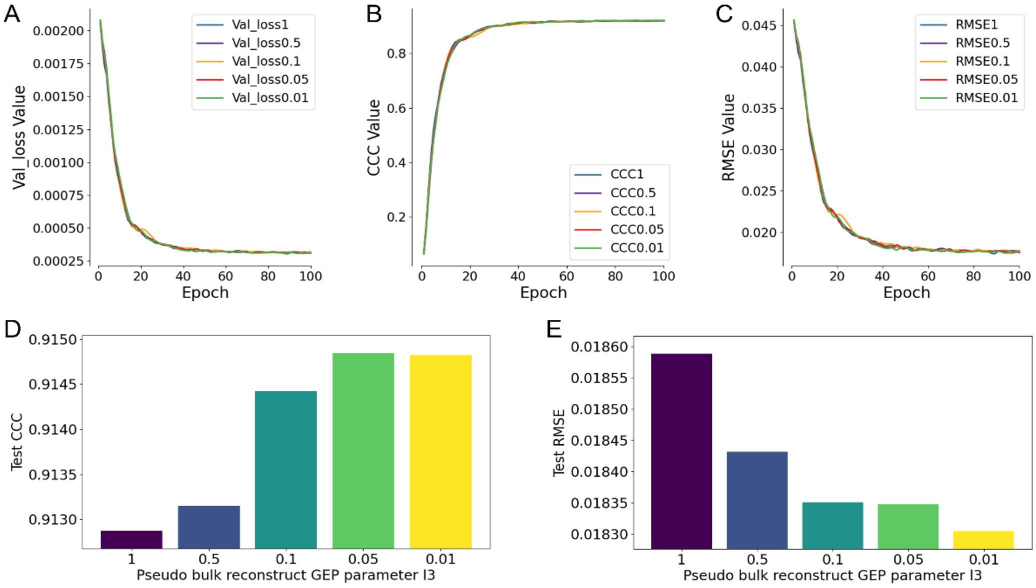

Supplementary Figure 10 Pseudo-bulk Reconstruction Loss (using predicted GEPs) Coefficient Optimization in Ascend Deconvolution Model (l3).

(A) Validation loss values across 100 epochs for different Pseudo-bulk reconstruction loss (using predicted GEPs) coefficients (λ): 1, 0.5, 0.1, 0.05, and 0.01. All the values of λ yield stable convergence throughout training with minimal differences from each other. (B) Concordance correlation coefficient (CCC) values on the validation set over epochs for different λ values. All the values of λ yield stable convergence of CCC throughout training with minimal differences from each other and they all converge to a high CCC value, demonstrating strong agreement between predicted and true values for all values of λ. (C) Root mean square error (RMSE) values on the validation set across epochs for each λ. All the values of λ yield stable convergence of RMSE throughout training with minimal differences from each other and they all converge to a low RMSE value which shows that all values of λ allow the model to minimize error effectively. (D) Test set CCC for each λ, showing that the smallest λ values (especially 0.05 and 0.01) produce the highest CCC values. Conversely, larger values (1) drastically reduce CCC, suggesting poorer predictive performance on the test set. (E) Test set RMSE for each λ, demonstrating that λ=0.01 achieves the lowest RMSE, followed by λ=0.05 and 0.1. Larger values (1) lead to substantial increases in RMSE, emphasizing that minimal Pseudo-bulk reconstruction loss (using predicted GEPs) constraint improves the model's accuracy. These results collectively highlight that a Pseudo-bulk reconstruction loss (using predicted GEPs) coefficient of 0.01 provides the best balance of low validation loss, high CCC, and low RMSE, supporting the selection of λ=0.01 for optimal performance in the deconvolution model.

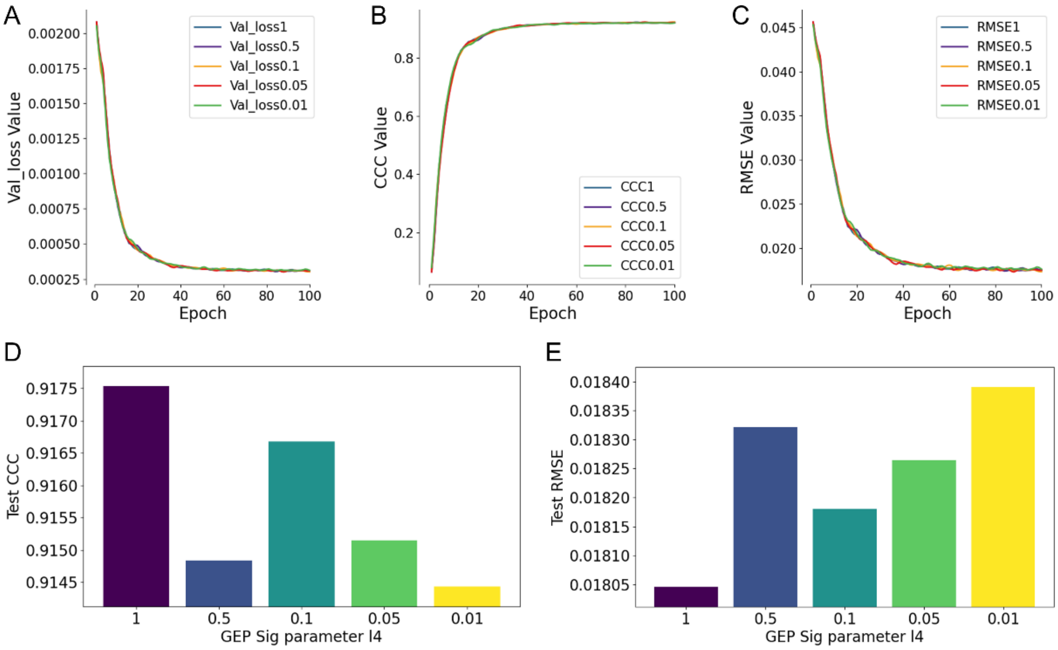

Supplementary Figure 11 Predicted GEP and Signature Matrix Loss Coefficient Optimization in Ascend Deconvolution Model (l4).

(A) Validation loss values across 100 epochs for different Predicted GEP and Signature Matrix Loss coefficients (λ): 1, 0.5, 0.1, 0.05, and 0.01. All the values of λ yield stable convergence throughout training with minimal differences from each other. (B) Concordance correlation coefficient (CCC) values on the validation set over epochs for different λ values. All the values of λ yield stable convergence of CCC throughout training with minimal differences from each other and they all converge to a high CCC value, demonstrating strong agreement between predicted and true values for all values of λ. (C) Root mean square error (RMSE) values on the validation set across epochs for each λ. All the values of λ yield stable convergence of RMSE throughout training with minimal differences from each other and they all converge to a low RMSE value which shows that all values of λ allow the model to minimize error effectively. (D) Test set CCC for each λ, showing that the highest λ value (1) produces the highest CCC values. Conversely, smaller values (0.01) drastically reduce CCC, suggesting poorer predictive performance on the test set. (E) Test set RMSE for each λ, demonstrating that λ=1 achieves the lowest RMSE. Smaller values (0.01) lead to substantial increases in RMSE, emphasizing that maximal Predicted GEP and Signature Matrix Loss constraint improves the model's accuracy. These results collectively highlight that a Predicted GEP and Signature Matrix Loss coefficient of 1 provides the best balance of low validation loss, high CCC, and low RMSE, supporting the selection of λ=1 for optimal performance in the deconvolution model.

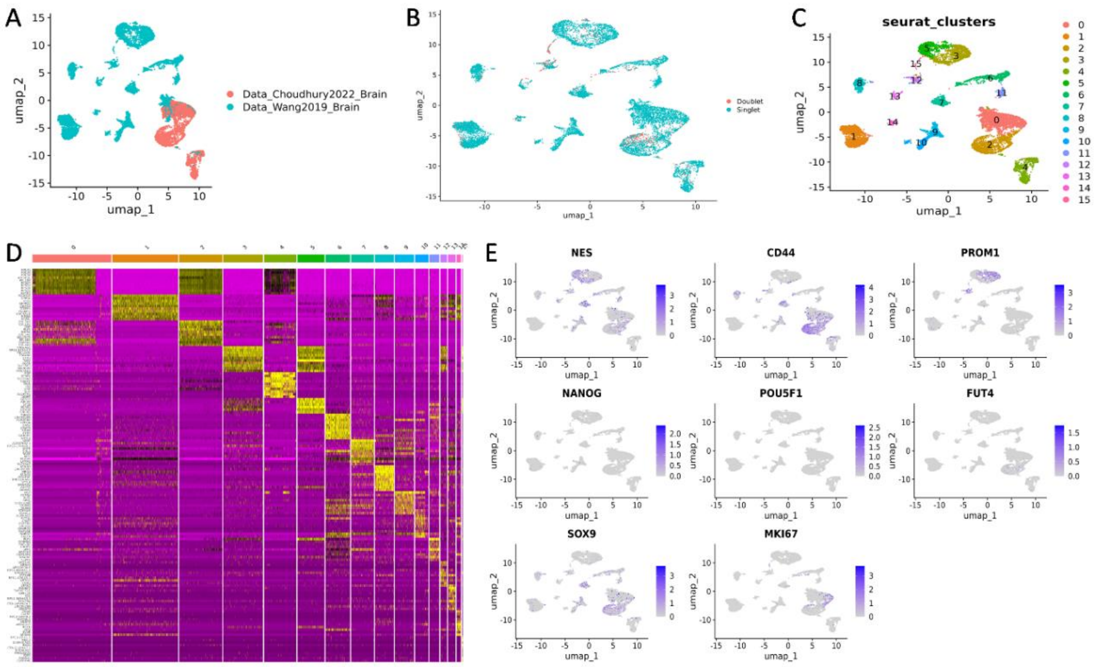

Supplementary Figure 12 Reannotation of malignant cells for identification of brain cancer stem cells.

Two different datasets were taken, and batch corrected for further processing (A). Mitochondrial expression-based data removal and doublet checking was done for data cleaning (B). Clusters were identified based on Louvain algorithm (C) and cluster-specific gene expression profile was obtained (D). Stem cell clusters were identified and then validated using marker expression profiles.

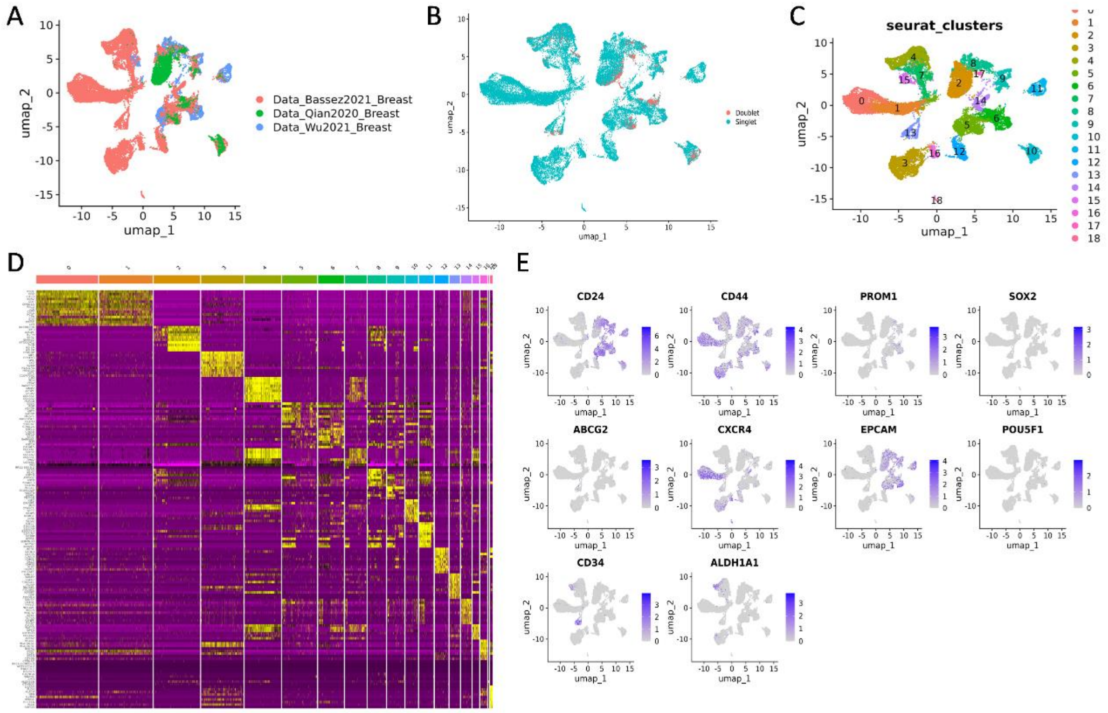

Supplementary Figure 13 Reannotation of malignant cells for identification of breast cancer stem cells.

Two different datasets were taken, and batch corrected for further processing (A). Mitochondrial expression-based data removal and doublet checking was done for data cleaning (B). Clusters were identified based on Louvain algorithm (C) and cluster-specific gene expression profile was obtained (D). Stem cell clusters were identified and then validated using marker expression profiles.

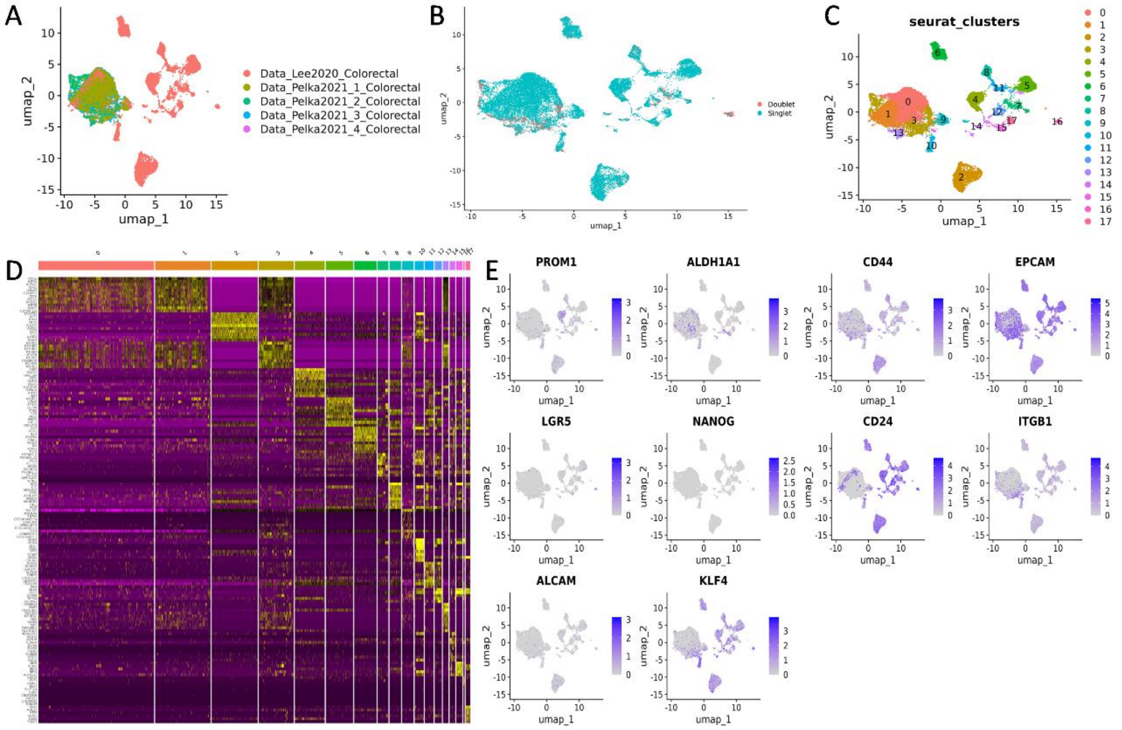

Supplementary Figure 14 Reannotation of malignant cells for identification of colorectal cancer stem cells.
Two different datasets were taken, and batch corrected for further processing (A). Mitochondrial expression-based data removal and doublet checking was done for data cleaning (B). Clusters were identified based on Louvain algorithm (C) and cluster-specific gene expression profile was obtained (D). Stem cell clusters were identified and then validated using marker expression profiles.

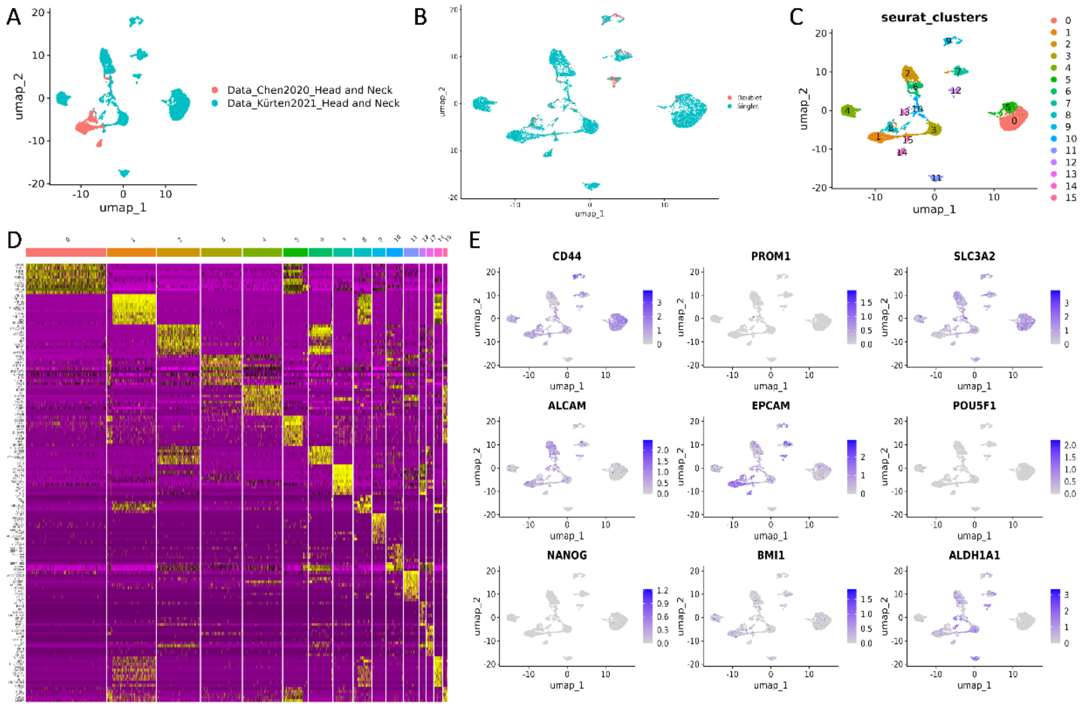

Supplementary Figure 15 Reannotation of malignant cells for identification of head and neck cancer stem cells.
Two different datasets were taken, and batch corrected for further processing (A). Mitochondrial expression-based data removal and doublet checking was done for data cleaning (B). Clusters were identified based on Louvain algorithm (C) and cluster-specific gene expression profile was obtained (D). Stem cell clusters were identified and then validated using marker expression profiles.

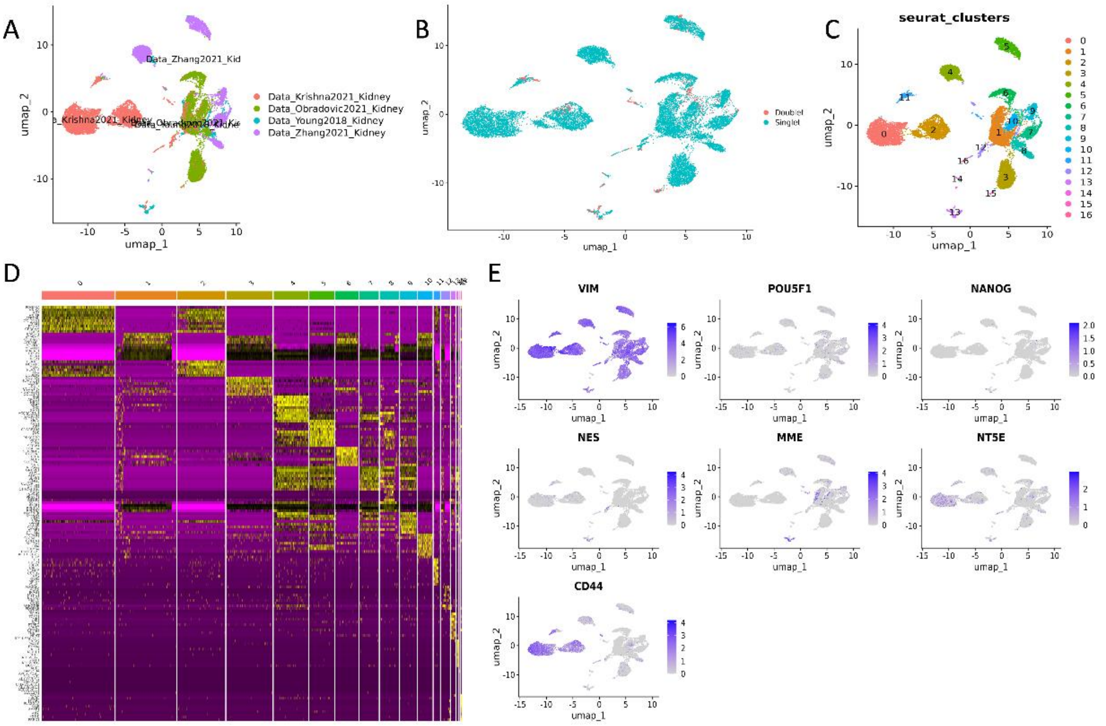

Supplementary Figure 16 Reannotation of malignant cells for identification of kidney cancer stem cells.
Two different datasets were taken, and batch corrected for further processing (A). Mitochondrial expression-based data removal and doublet checking was done for data cleaning (B). Clusters were identified based on Louvain algorithm (C) and cluster-specific gene expression profile was obtained (D). Stem cell clusters were identified and then validated using marker expression profiles.

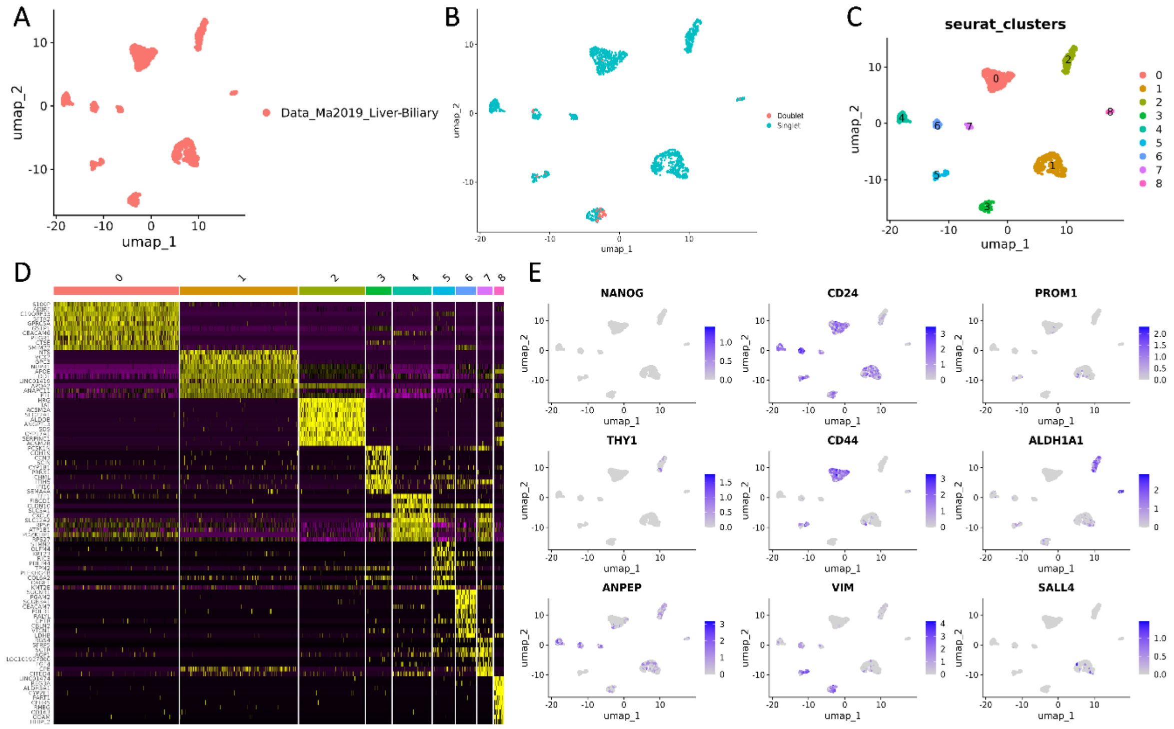

Supplementary Figure 17 Reannotation of malignant cells for identification of liver cancer stem cells.
Two different datasets were taken, and batch corrected for further processing (A). Mitochondrial expression-based data removal and doublet checking was done for data cleaning (B). Clusters were identified based on Louvain algorithm (C) and cluster-specific gene expression profile was obtained (D). Stem cell clusters were identified and then validated using marker expression profiles.

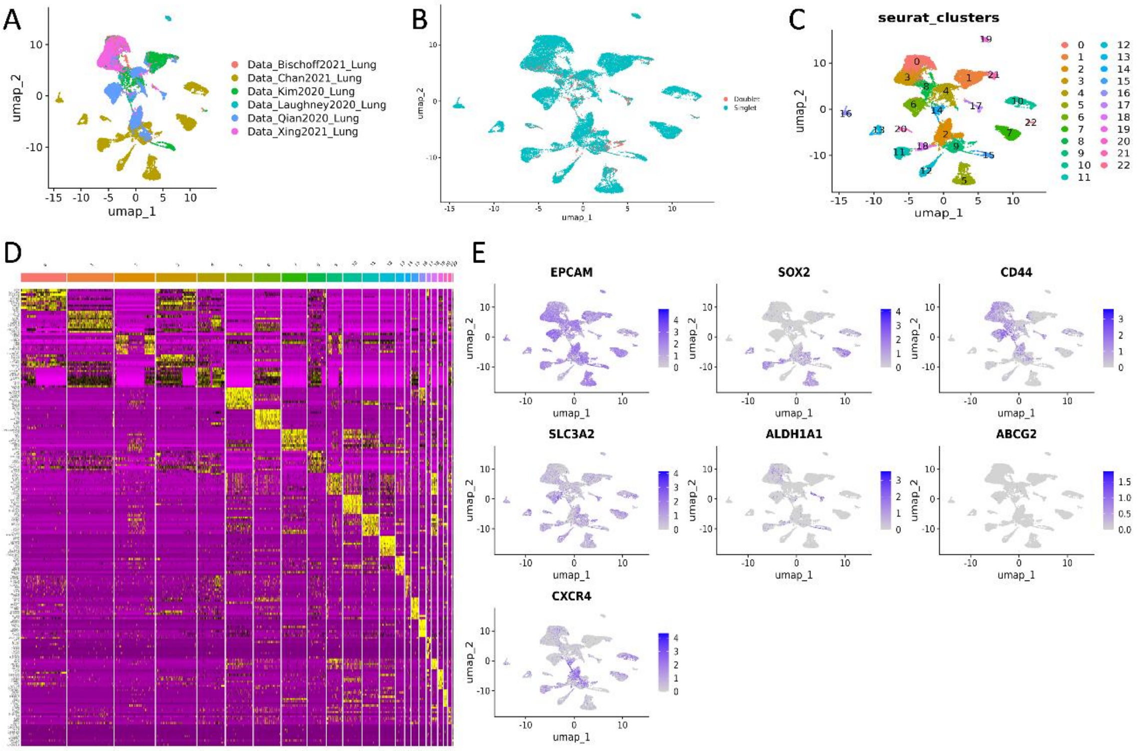

Supplementary Figure 18 Reannotation of malignant cells for identification of lung cancer stem cells.
Two different datasets were taken, and batch corrected for further processing (A). Mitochondrial expression-based data removal and doublet checking was done for data cleaning (B). Clusters were identified based on Louvain algorithm (C) and cluster-specific gene expression profile was obtained (D). Stem cell clusters were identified and then validated using marker expression profiles.

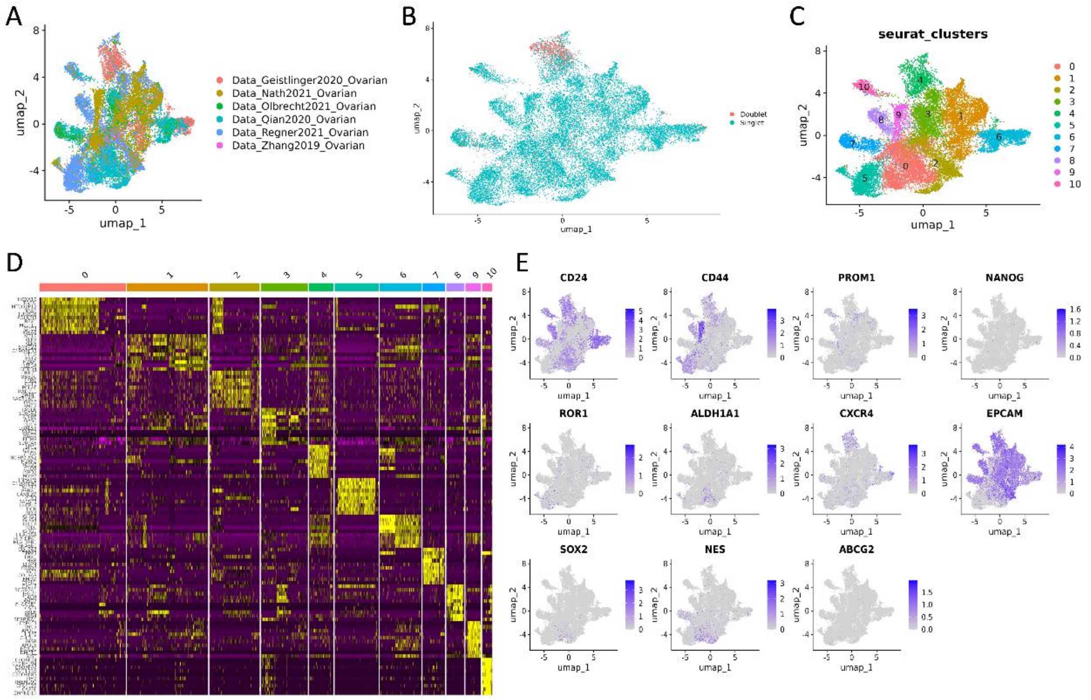

Supplementary Figure 19 Reannotation of malignant cells for identification of ovarian cancer stem cells.
Two different datasets were taken, and batch corrected for further processing (A). Mitochondrial expression-based data removal and doublet checking was done for data cleaning (B). Clusters were identified based on Louvain algorithm (C) and cluster-specific gene expression profile was obtained (D). Stem cell clusters were identified and then validated using marker expression profiles.

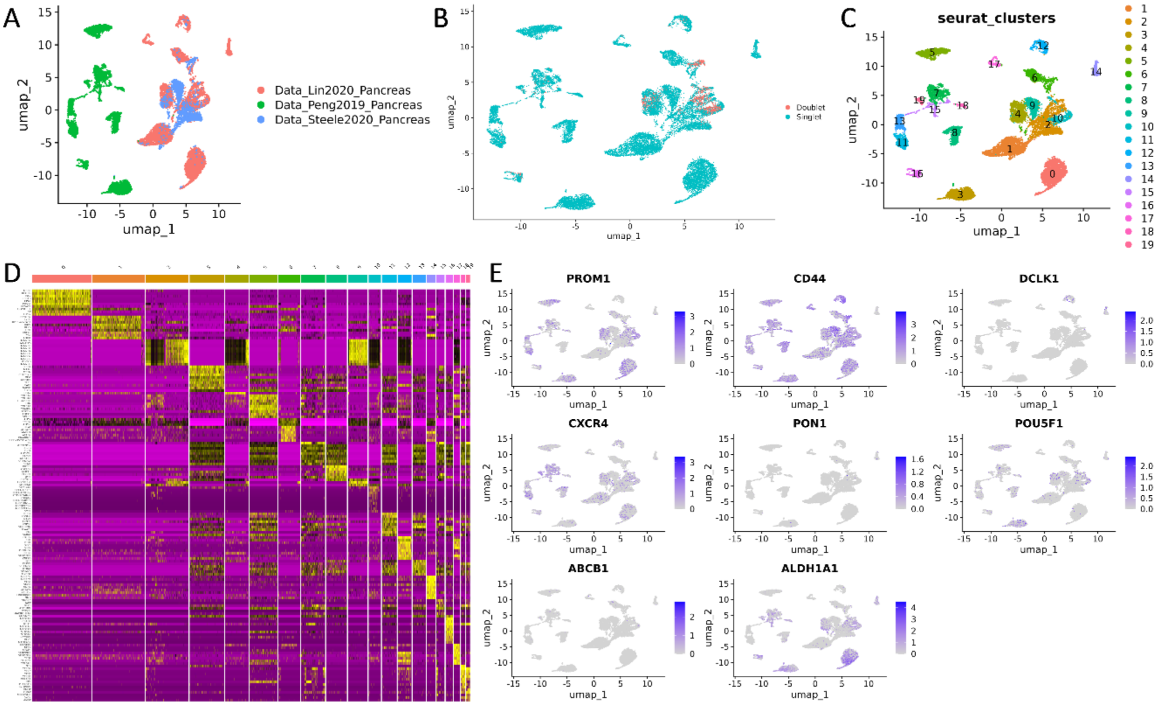

Supplementary Figure 20 Reannotation of malignant cells for identification of pancreatic cancer stem cells. Two different datasets were taken, and batch corrected for further processing (A). Mitochondrial expression-based data removal and doublet checking was done for data cleaning (B). Clusters were identified based on Louvain algorithm (C) and cluster-specific gene expression profile was obtained (D). Stem cell clusters were identified and then validated using marker expression profiles.

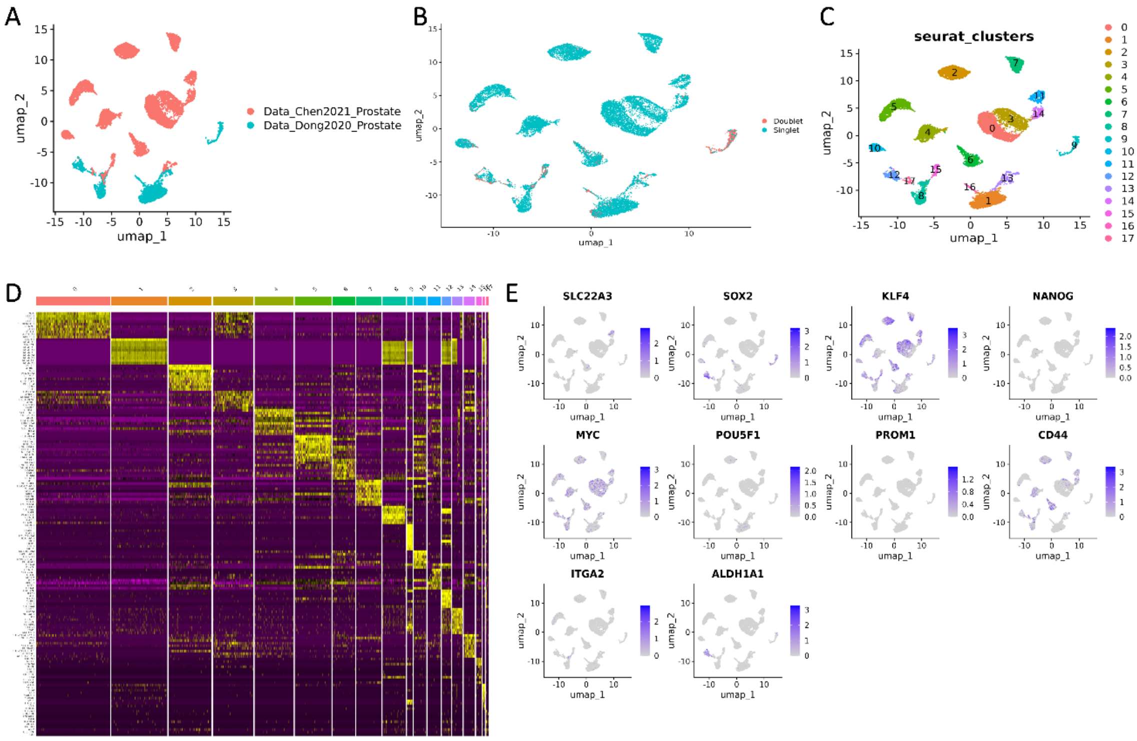

Supplementary Figure 21 Reannotation of malignant cells for identification of prostate cancer stem cells. Two different datasets were taken, and batch corrected for further processing (A). Mitochondrial expression-based data removal and doublet checking was done for data cleaning (B). Clusters were identified based on Louvain algorithm (C) and cluster-specific gene expression profile was obtained (D). Stem cell clusters were identified and then validated using marker expression profiles.

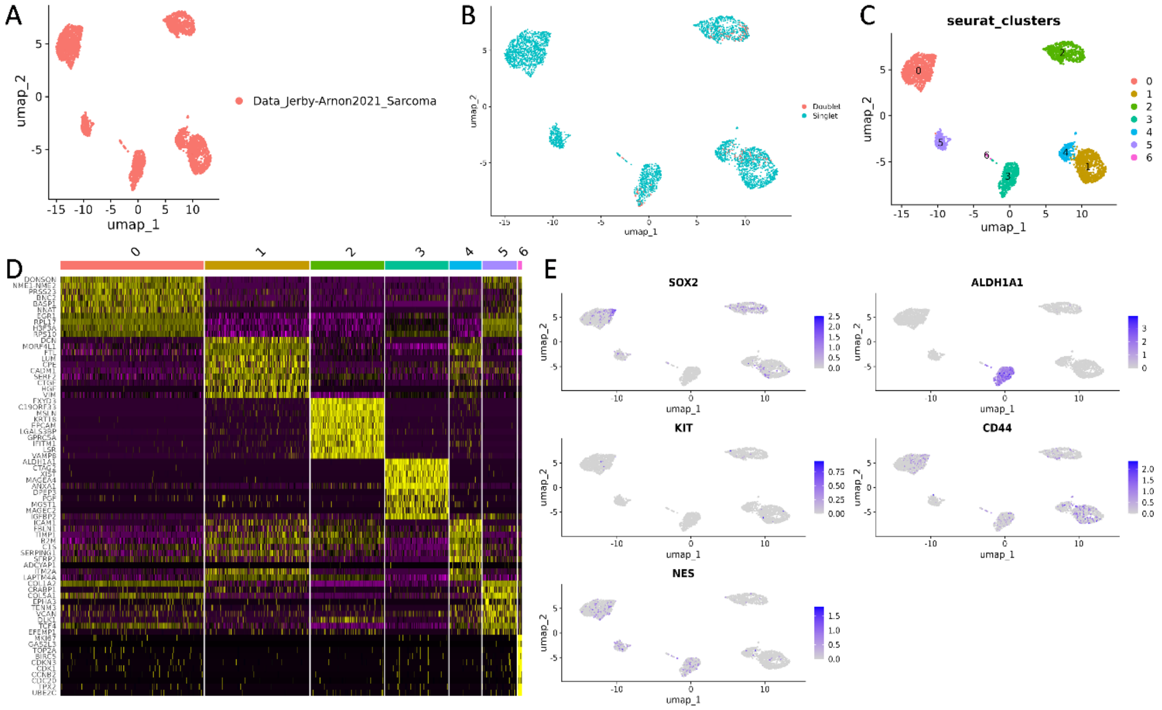

Supplementary Figure 22 Reannotation of malignant cells for identification of sarcoma cancer stem cells.
Two different datasets were taken, and batch corrected for further processing (A). Mitochondrial expression-based data removal and doublet checking was done for data cleaning (B). Clusters were identified based on Louvain algorithm (C) and cluster-specific gene expression profile was obtained (D). Stem cell clusters were identified and then validated using marker expression profiles.

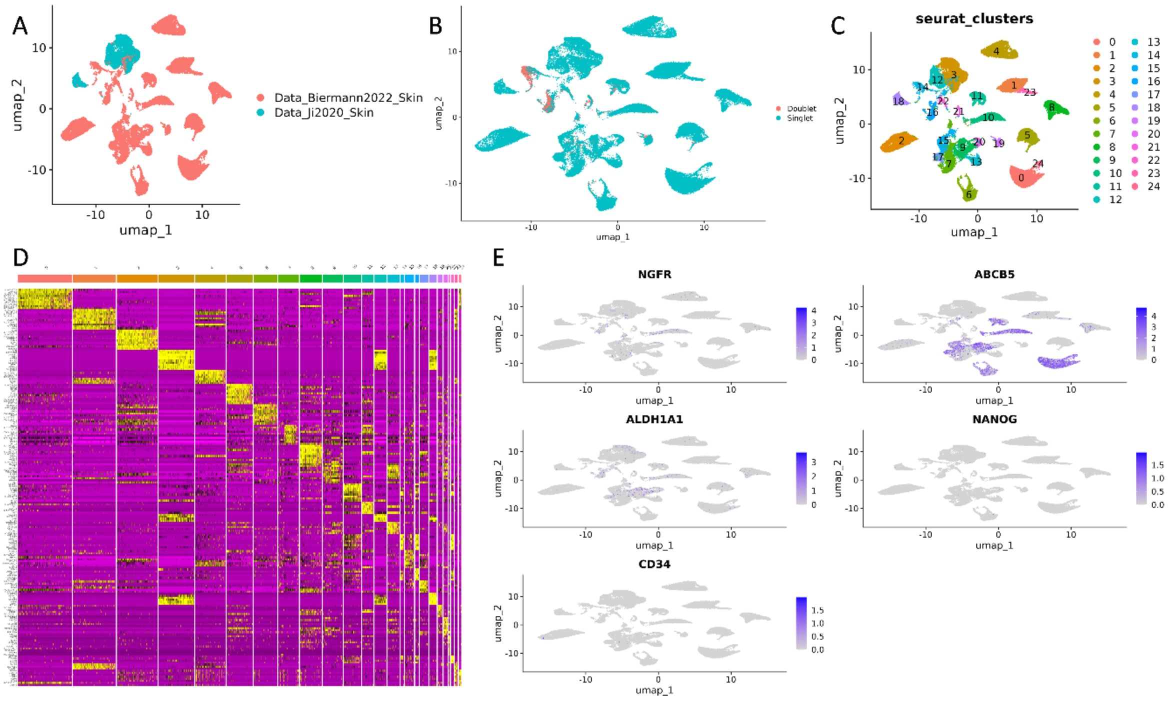

Supplementary Figure 23 Reannotation of malignant cells for identification of skin cancer stem cells.
Two different datasets were taken, and batch corrected for further processing (A). Mitochondrial expression-based data removal and doublet checking was done for data cleaning (B). Clusters were identified based on Louvain algorithm (C) and cluster-specific gene expression profile was obtained (D). Stem cell clusters were identified and then validated using marker expression profiles.

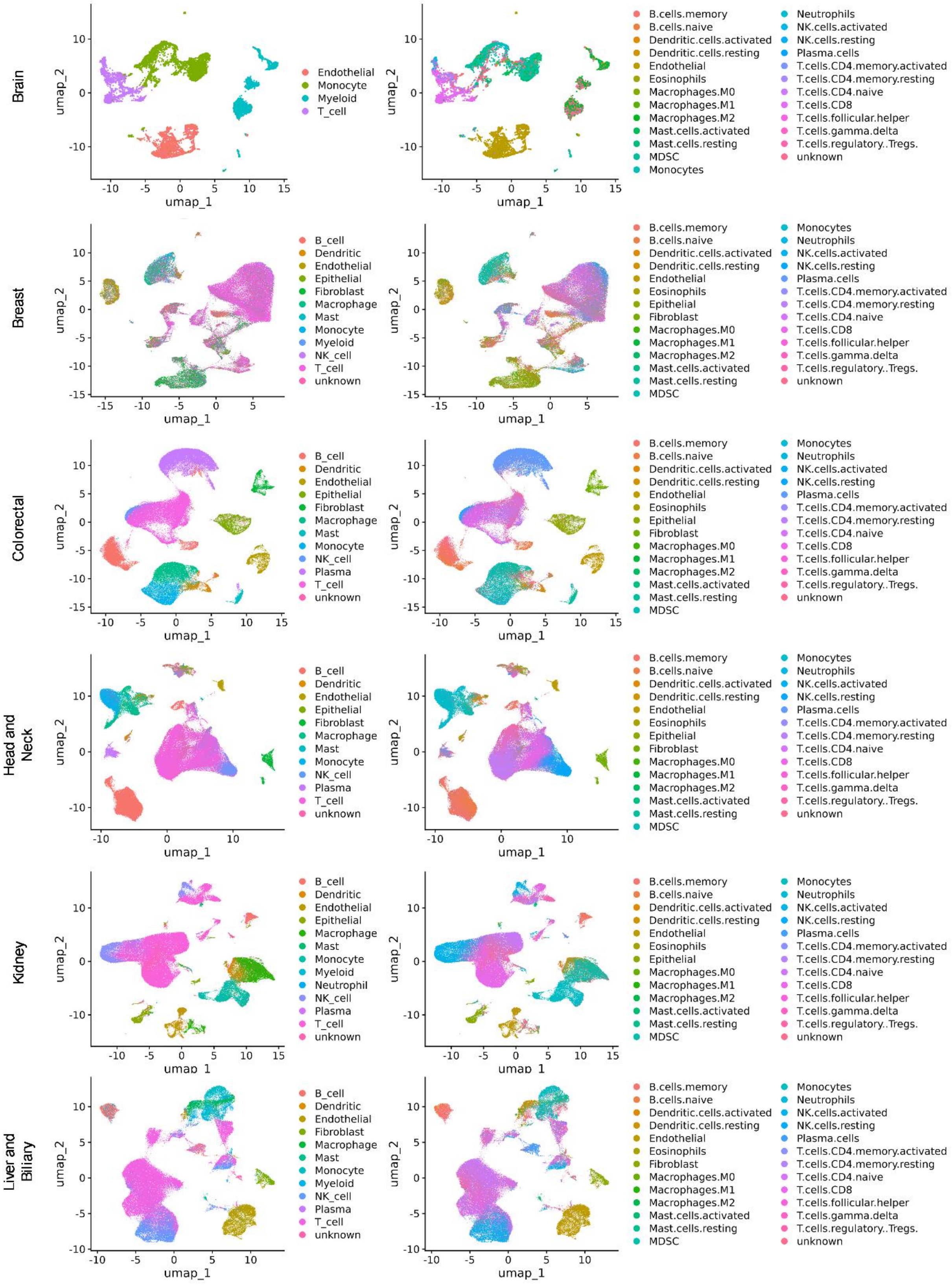

Figure continued… next page……

Figure legend and descriptor: next page

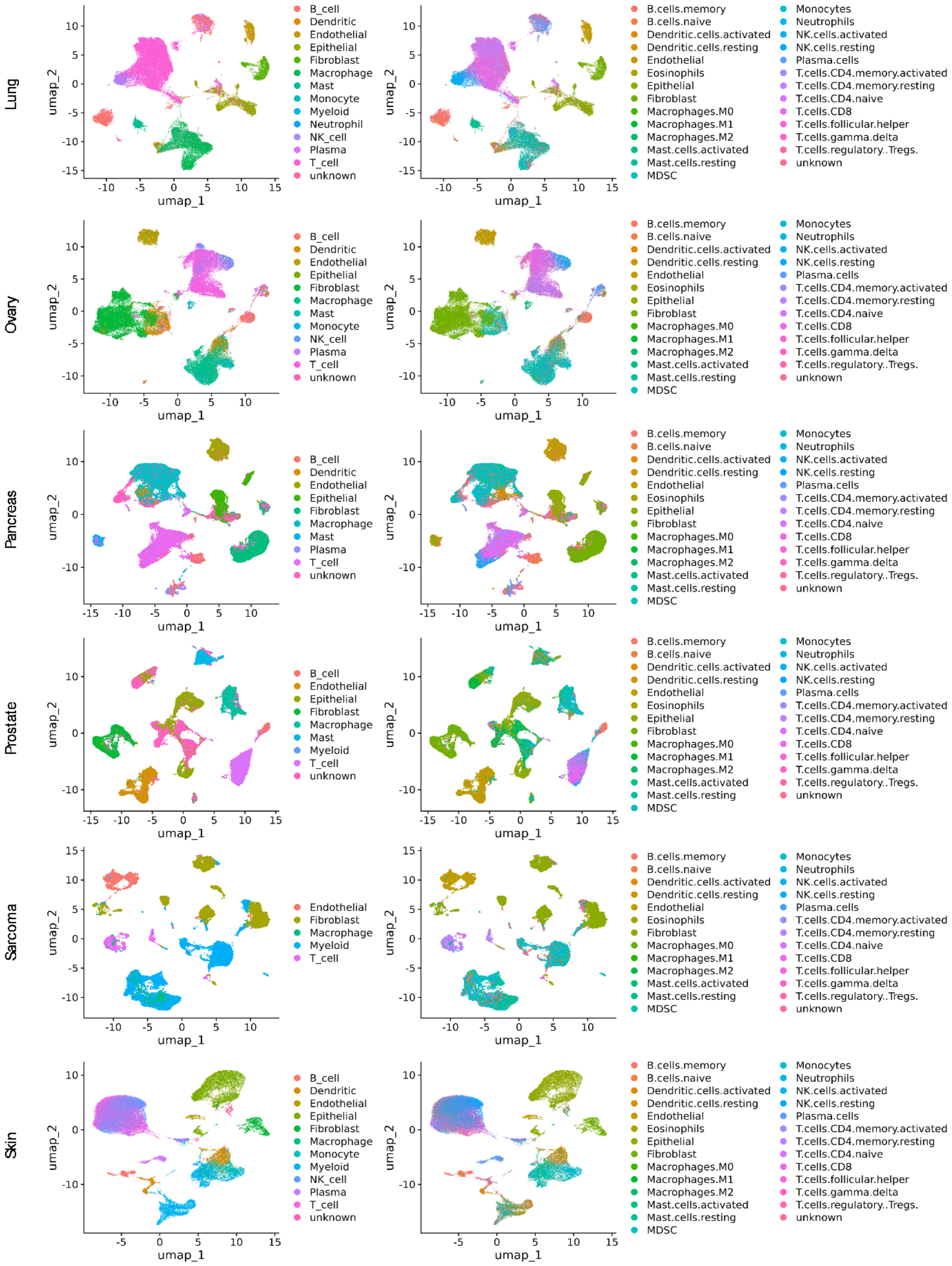

Supplementary Figure 24 Reannotation of cancer-specific immune infiltrates.
The left panel shows the batch corrected UMAP of tissue-specific merged annotations of immune cell types collected from tumor scRNAseq data sources. For each tumor tissue, the right panel shows the UMAP representation of reannotated cell types. Common cell types before, and after reannotation shows similar representation confirming the robustness.

Supplementary Table 1 Hyperparameters for all validation datasets

| | PBMC | Development | Malignant | Mouse data | Malignant-test |
|---|---|---|---|---|---|
| Loss coefficients | | | | | |
| L1 | 0.01 | 0.01 | 0.01 | 0.01 | 0.01 |
| L2 | 0.01 | 0.01 | 0.01 | 0.01 | 0.005 |
| L3 | 0.01 | 0.01 | 0.01 | 0.01 | 0.005 |
| L4 | 1.0 | 1.0 | 1.0 | 1.0 | 0.1 |
| Learning rate | 0.0001 | 0.0001 | 0.0001 | 0.0001 | 0.0001 |
| Dropout | 0.2 | 0.1 | 0.1 | 0.1 | 0.1 |
| Latent dimension | 512 | 512 | 512 | 512 | 512 |

Supplementary Table 2 List of curated CSC markers

| Primary tumor tissue | CSC markers | References |
|---|---|---|
| Brain | NES, CD44, PROM1, NANOG, SOX2, POU5F1, FUT4, ITGB6, SOX9, MKI67, ALDH1A1 | 1–4 |
| Breast | CD44, PROM1, SOX2, ABCG2, CXCR4, EPCAM, POU5F1, CD34, ALDH1A1 | 5–8, 9 |
| Colorectum | PROM1, ALDH1A1, CD44, EPCAM, LGR5, NANOG, CD24, ITGB1, ALCAM, KLF4 | 10–13 |
| Head & Neck | CD44, PROM1, SLC3A2, ALCAM, EPCAM, POU5F1, NANOG, BMI1, ALDH1A1 | 14–19 |
| Kidney | VIM, POU5F1, NANOG, NES, MME, CD29, NT5E, CD44, CXCR4 | 20–22 |
| Liver | NANOG, CD24, PROM1, THY1, CD44, ICAM1, EPCAM, ALDH1A1, ANPEP, VIM, SALL4 | 23–26 |
| Lung | EPCAM, SOX2, CD44, SLC3A2, ALDH1A1, ABCG2, CXCR4, MSI2 | 27–32 |
| Ovary | CD24, CD44, PROM1, NANOG, ROR1, ALDH1A1, CXCR4, VCAM1, EPCAM, SOX2, NES, ABCG2 | 33–36,36–40 |
| Pancreas | PROM1, CD24, CD44, DCLK1, CXCR4, PON1, POU5F1, ABCB1, ALDH1A1 | 41–45 |
| Prostate | SLC22A3, SOX2, KLF4, NANOG, MYC, POU5F1, PROM1, CD44, ALDH1A1 | 46–49 |
| Sarcoma | SOX2, ALDH1A1, KIT, PROM1, STRO1, ENG, CD44, POU5F1, NANOG, NES, CD24 | 50–53 |
| Skin | NGFR, ABCB5, ALDH1A1, NANOG, CD34 | 54–60 |

### Supplementary Methods

#### Data curation for stem cell classifier training:

In our classification of stem cells based on differentiation potential, we have identified three main categories: pluripotent, multipotent, and unipotent stem cells[61–63]. This classification enhances our understanding of their unique properties and functions, which are essential for applications in understanding stem cell heterogeneity and differentiation/dedifferentiation path.

Pluripotent Stem Cells are characterized by their ability to differentiate into nearly all cell types in the body[64]. Embryonic stem cells (ESCs) are the primary example of this category, derived from the inner cell mass of the blastocyst[65,66]. Human embryonic stem cells (hESCs) can exist in two states: naive and primed[67,68]. Naive hESCs resemble the inner cell mass of the blastocyst, exhibiting characteristics associated with early developmental stages, such as high expression of pluripotency markers. In contrast, primed hESCs are developmentally more advanced, similar to the post-implantation epiblast, and are considered to be in a primed pluripotent state. Both naive and primed populations are mostly homogeneous with no clear lineage-related structure and go through an intermediate subpopulation of naive cells with primed-like expression[69]. Therefore, capturing both naive and primed human embryonic stem cells (hESCs) is essential to fully define pluripotency at the transcriptomic level. Including a dataset with both naive and primed hESCs allows us to combine their unique transcriptomic profiles, thereby enhancing our understanding of pluripotency in humans.

In the multipotent category, mesenchymal stem cells were chosen for their capacity to differentiate into various connective tissues like bone, cartilage, and fat[70–72]. Neural stem cells were included because they can generate neurons and glial cells, playing a vital role in brain repair and neurogenesis[73,74]. Neural crest cells were selected due to their versatility in forming diverse cell types, including neurons, glial cells, and pigment cells, highlighting their importance in developmental biology[75–78]. Skin stem cells were included for their role in the constant renewal and repair of the skin, demonstrating their significance in maintaining tissue integrity and healing[79–82].

Oligopotency and unipotency represent closely related stages in the functional hierarchy of adult stem cells, forming a continuum of differentiation potential. Oligopotent stem cells can generate a few closely related cell types, while unipotent stem cells are restricted to producing only one specific cell type. This relationship is evident in systems like hematopoiesis, where multipotent stem cells transition through oligopotent stages (e.g., lymphoid progenitors) before becoming unipotent precursors[83]. Similarly, adult epithelial and germline stem cells illustrate this continuum by progressing through increasingly restricted potency states based on tissue needs and regenerative requirements. This functional overlap is crucial for somatic tissues, where strictly unipotent states, such as germ stem cells, are less relevant due to the broader repair and regenerative needs of diseases affecting somatic tissues, such as somatic cell cancers. For categorizing the lowest potency state of CSCs, their close functional relationship supports classifying oligopotent and unipotent states together, particularly when higher potency states like pluripotency and multipotency are already distinguished. These stages highlight the practical continuum in stem cell differentiation that supports tissue-specific regenerative capacity while remaining responsive to somatic conditions.

In the oligopotent/unipotent category, oligopotent motor neuron progenitors were selected. In experimental conditions, motor neuron progenitors can sometimes be reprogrammed to adopt other fates through genetic manipulation or exposure to novel signals. However, under normal physiological conditions, they remain of limited potency, producing only subventricular motor neurons, when only subventricular motor neuron progenitors are taken[84]. Satellite cells were included due to their capacity to produce only myogenic precursor cells in-vivo, playing an essential role in muscle maintenance and repair after injury[85,86], despite its controversial differentiation into multiple lineages in-vitro. Alveolar type II cells were selected for their function in pulmonary surfactant production and their role in regenerating alveolar epithelial cells following lung injury, underscoring their importance in respiratory health[87–89]. AT2 cells act as progenitor cells in the alveoli. They proliferate and differentiate into AT1 cells to repair alveolar damage, ensuring the integrity of the epithelial barrier. This lineage commitment to AT1 cells underlines their unipotent nature. It is important to note that, AT2 and AT1 both have a common bipotent progenitor, which we have not considered here. We have only considered unipotent AT2 cells, for our analysis. This classification facilitates understanding their unique functions and therapeutic potential. Including totipotent stem cells was not necessary, as they are only present in the early stages of development, with no totipotent cells remaining after blastulation, limiting their relevance in later developmental and clinical contexts[66].

Classifiers often classify data into predetermined categories, even in the absence of those categories in the ground truth data. This limitation is evident in tools like CytoTRACE or CytoTRACE 2, which may erroneously classify cells as totipotent or differentiated despite the input data lacking these specific cell types, potentially leading to misleading biological interpretations[90,91]. Moreover, qualitative classifications by these models consider cooccurrence of totipotent and differentiated cells. Developmentally totipotent and differentiated cells cannot co-occur, as totipotent cells have the unique ability to give rise to all cell types, including extraembryonic tissues, while differentiated cells have committed to specific lineages. While totipotent stem cells can give rise to all cell types, including extra-embryonic tissues, they are not included in this classification since they are only present in the very early stages of development[92]. After the blastulation stage, no totipotent cells remain, making their consideration less relevant for most applications.

##### Comparison of data imbalance handling approaches for Stem cell classifier module:

To tackle the significant class imbalance in our dataset, where Multipotent samples were approximately eight times more abundant than Pluripotent and Unipotent samples, we designed a three-stage training process. Each stage utilized a different strategy to balance the data, enabling us to assess model performance across various scenarios and ultimately select the optimal approach for our final classifier.

1. Stage 1: Baseline Model Training on Imbalanced Data
   In the first stage, we trained a set of initial models on the unaltered imbalanced dataset, containing Multipotent, Pluripotent, and Unipotent samples in their original proportions. This setup allowed us to establish a baseline performance without any interventions to address the imbalance. While this imbalanced data provided insights into the models' natural performance tendencies, the results revealed a pronounced bias toward the majority class (Multipotent), with noticeably lower accuracy and recall for the minority classes (Pluripotent and Unipotent). These results underscored the need for additional strategies to mitigate class imbalance and improve classification for underrepresented classes.

2. Stage 2: Model Training with Synthetic Minority Over-sampling Technique (SMOTE)[93,94]
   To address these issues, we applied SMOTE in the second stage to create a balanced dataset. SMOTE generated synthetic samples for the minority classes, Pluripotent and Unipotent, resulting in a dataset with nearly equal representation across all classes. Training the models on this SMOTE-adjusted dataset improved classification performance for Pluripotent and Unipotent classes, reducing bias toward the majority class. However, balancing the dataset in this manner introduced some concerns, such as the risk of overfitting due to synthetic data, which may not capture the true variability of the original samples. Although the SMOTE approach helped mitigate imbalance effects, the potential for model generalization issues prompted us to explore alternative balancing techniques.

3. Stage 3: Ensemble Approach with Balanced Representation
   In the final stage, we implemented an ensemble-based approach to balance the data without generating synthetic samples[95,96]. We divided the Multipotent samples into eight distinct subsets, retaining all Pluripotent and Unipotent samples in each subset. For each model in the ensemble, we trained on a unique subset of Multipotent samples combined with the complete sets of Pluripotent and Unipotent samples. This method ensured balanced representation across the models in the ensemble, addressing class imbalance while preserving the original sample distribution. By using this technique, we avoided both the limitations of the unaltered imbalance in Stage 1 and the potential overfitting risks associated with synthetic data in Stage 2.

After training the models across all three stages, we evaluated their performance using metrics such as accuracy, precision, recall, and F1 score, focusing on balanced performance across all classes. Cross-validation results showed that the ensemble-based models from Stage 3 significantly outperformed the models trained on both the imbalanced dataset (Stage 1) and the SMOTE-balanced dataset (Stage 2). Specifically, the Logistic Regression (LR) model in the ensemble achieved the highest accuracy, precision, and recall across the majority and minority classes, demonstrating a well-rounded and consistent performance. Consequently, we selected the LR model from the ensemble approach as the optimal classifier for our final implementation. Interestingly, the Logistic Regression model exhibited nearly identical performance on both the imbalanced dataset (Stage 1) and the ensemble-based dataset (Stage 3). This unusual consistency suggests that LR's inherent ability to handle imbalance effectively may have played a role. Logistic Regression tends to perform well in binary and multiclass settings with imbalanced data due to its probabilistic nature, which can create stable decision boundaries without being heavily skewed by class proportions. However, in the ensemble approach, the partitioning of the data likely further refined LR’s decision boundaries, allowing it to generalize even more effectively across minority classes without losing performance on the majority class. The combination of LR’s robustness to imbalance and the balanced representation in the ensemble approach created a synergy, resulting in high, uniform performance across all evaluation metrics.

##### scRNAseq data preparation for tumor deconvolution model training:

The single-cell RNA sequencing (scRNA-seq) gene expression matrix for various cancer types was obtained from the publicly available Weizmann Institute's 3CA dataset, which contains high-resolution data on cancer-associated cell types[97]. The raw data were preprocessed and subjected to stringent quality control using the Seurat package (v5.1.0) in R, a widely used toolkit for the analysis of scRNA-seq data[98,99]. Initial filtering was performed to retain only high-quality cells. Cells that expressed fewer than two hundred genes were excluded to avoid the inclusion of empty droplets or low-quality cells, while cells expressing more than 6,000 genes were removed as potential doublets or artifacts. Additionally, cells with a mitochondrial gene fraction exceeding 10% were eliminated, as an elevated fraction of mitochondrial RNA typically indicates dying or stressed cells, which can skew downstream analyses. To further remove potential doublets—artifacts where two or more cells are captured together, leading to an artificially high gene expression profile—the DoubletFinder package (v2.0) in R was employed[100]. Doublet detection was performed by first determining the optimal pK parameter, which controls the neighborhood size used to estimate the probability of doublet formation. This was achieved using the paramSweep function, which iterates through a range of pK values to identify the most suitable one for the dataset. The BCmetric values obtained from the paramSweep results were summarized using the find.pK function, and the pK value corresponding to the maximum BCmetric was selected as the optimal parameter for detecting doublets. Since homotypic doublets (doublets formed from similar cell types) are more difficult to detect than heterotypic doublets, adjustments were made to account for their presence. The expected doublet rate was corrected based on the proportion of homotypic cells, adjusting the nExp_poi.adj parameter accordingly. This adjustment helps in more accurately identifying and filtering out homotypic doublets. After determining the expected number of doublets and performing the necessary adjustments, doublets were classified and removed from the dataset. Only the remaining high-confidence singlets were retained for downstream analyses, such as clustering, differential expression analysis, and trajectory inference, ensuring the integrity of the biological insights derived from the data. This preprocessing pipeline, combining Seurat for general quality control and DoubletFinder for doublet identification, ensures that the final dataset is clean and reliable, with minimal contamination from low-quality cells and doublets.

##### Immune Annotation:

Cell type annotations for major non-immune populations, including malignant cells, fibroblasts, epithelial cells, and endothelial cells, were directly preserved from the original dataset provided by the Weizmann Institute’s 3CA. These annotations were initially assigned based on gene expression profiles characteristic of each cell type and validated through lineage-specific markers included in the dataset’s metadata. However, for the detailed classification of immune cell populations, more specialized tools were required due to the complexity of immune infiltrates in the tumor microenvironment (TME). To achieve robust immune cell annotations, the SCINA (Semi-supervised Category Identification and Assignment) package was employed, leveraging known marker genes to classify immune cell populations[101]. SCINA is designed for supervised annotation of scRNA-seq data, using a set of predefined marker genes. For this analysis, the LM22 matrix, derived from the study by Newman et al., served as the primary reference for immune cell markers[102]. The LM22 matrix is a widely recognized set of gene markers used to deconvolute immune cell types in both bulk and single-cell RNA sequencing data, encompassing major immune populations such as T cells, B cells, macrophages, and natural killer (NK) cells. However, the LM22 matrix does not include marker genes for myeloid-derived suppressor cells (MDSCs), which play a crucial role in cancer progression by modulating immune responses and fostering a pro-tumorigenic microenvironment. Given the importance of accurately identifying MDSCs in the TME, a separate set of MDSC marker genes was curated from external study. Specifically, these markers were obtained from a key study on MDSCs, which detailed the molecular characteristics of these cells in cancer[103]. Following the inclusion of MDSC markers, SCINA was run using the combined marker gene set—both from LM22 and the custom MDSC gene list. This process allowed for the precise annotation of immune cells, including major populations such as CD4+ and CD8+ T cells, B cells, macrophages, NK cells, dendritic cells, and MDSCs, thereby providing a comprehensive view of the immune landscape within the tumor microenvironment.

##### Cancer Stem Cells annotation:

The annotation of cancer stem cells (CSCs) was carried out using a multi-step, manual approach designed to maximize precision, particularly given the challenges in distinguishing CSCs from normal stem cells. This process began with the isolation of malignant cells, leveraging the original annotations and post-quality control filtering of the dataset. The key distinction here is that stem cell annotation was restricted to malignant cells only, as cancerous tissues often contain both normal stem cells and CSCs, and it was crucial to differentiate the two. This approach improves upon existing tools like CytoTRACE, which cannot accurately distinguish between normal and cancer stem cells, especially in a mixed population. By focusing solely on malignant cells, our method ensures a more robust and cancer-specific identification of CSCs. The gene expression matrix of malignant cells was first preprocessed using the Seurat package (v5.1.0) in R. The NormalizeData function was employed to normalize the data, using the LogNormalize method, which scales gene expression levels by a factor of 10,000 and subsequently log-transforms the data. This standardizes the gene expression across all cells, making comparisons more reliable. Next, the FindVariableFeatures function was used to detect the top 2,000 highly variable genes across cells, employing the variance-stabilizing transformation (vst) method to capture the most biologically relevant genes with the highest variation across the dataset. This step is critical in scRNA-seq analysis, as it ensures that the most informative genes are prioritized for downstream analyses. The data were then scaled using ScaleData, which standardizes the expression levels for each gene, thereby reducing potential bias caused by overrepresented or underrepresented genes in the dataset. To reduce the dimensionality of the dataset and facilitate efficient clustering, Principal Component Analysis (PCA) was performed using the RunPCA function, focusing on the top 2,000 highly variable genes. The first twenty principal components (PCs) were retained for further analysis, as these PCs capture most of the variance within the dataset. However, since the dataset was derived from multiple experimental conditions, it was necessary to address potential batch effects that could obscure biological signals. This was achieved through the RunHarmony function from the Harmony package[104], which corrects for technical variation while preserving biological differences across cells. Harmony aligns cells from different batches into a shared space, enabling more accurate clustering and cell-type identification. Following batch correction, FindNeighbors was used to construct a nearest neighbor graph, which forms the basis for clustering cells into distinct groups. The Louvain algorithm, implemented in Seurat's FindClusters function, was then applied to partition cells into clusters. The resolution parameter was set to 0.3 to ensure fine-grained clustering of cell populations. After clustering, violin plots were generated to visualize the expression levels of known cancer stem cell markers for each cancer type across the clusters, providing a qualitative assessment of potential CSC-like clusters. To quantitatively assess stem cell phenotypes, UCell analysis was performed on each cluster using cancer type-specific stem cell marker genes. UCell is a robust single-cell enrichment analysis method that calculates a score for each cell based on the expression of a predefined set of marker genes[105]. In this case, stem cell marker genes tailored to specific cancer types were used, and the UCell scores were calculated for all clusters. Clusters exhibiting high UCell scores, and elevated expression of these stem cell markers were identified as potential CSC clusters.

#### Pseudobulk data generation

##### PBMC pseudobulk data generation

To process and annotate single-cell RNA sequencing (scRNA-seq) data, we utilized publicly available data from the 10x Genomics platform comprising 20,000 peripheral blood mononuclear cells (PBMCs). The raw data were accessed from the 10x Genomics filtered_feature_bc_matrix.h5 file using the Seurat package (v4.0) in R. The data were first loaded using the Read10X_h5 function, and a Seurat object was created with the function CreateSeuratObject(). Cells were filtered based on key quality control metrics: cells with fewer than 800 RNA counts, fewer than five hundred detected features, or mitochondrial gene content exceeding 10% were excluded. This yielded a high-quality PBMC dataset for downstream analysis. We then normalized the data using the NormalizeData() function in Seurat, followed by the identification of highly variable features (FindVariableFeatures()). The data were scaled using ScaleData(), and principal component analysis (PCA) was conducted on the top 20 principal components (RunPCA()). Nearest-neighbor graphs were constructed with the FindNeighbors() function, and clustering was performed using the Louvain algorithm (FindClusters()). To visualize the clustering results, Uniform Manifold Approximation and Projection (UMAP) was applied (RunUMAP()), and the resulting UMAP plot was used to display the spatial distribution of cell clusters.

For cell-type annotation, we employed SingleR (v1.4)[106] in conjunction with the Human Primary Cell Atlas reference dataset from the celldex R package. Expression data from the PBMC Seurat object were extracted using GetAssayData() and passed as the test dataset to SingleR() for automatic annotation. SingleR compared the PBMC expression profiles to the reference, assigning each cell a predicted label based on its similarity to primary cell types. The resulting cell-type labels were appended to the Seurat metadata and visualized on the UMAP plot. To assess the confidence of the annotation, we extracted the prediction scores from SingleR.

To generate training data, we prepared pseudobulk data using SimBu[107]. We began by creating a dataset using the dataset() function, providing filtered annotations, count matrix, and TPM matrix. We used the simulate_bulk() function to simulate pseudobulk data, specifying scenario = "random" to introduce random variation. Additionally, we set ncells = 100 to select 100 cells per pseudobulk sample, and nsamples = 1000 to create 1000 bulk samples. We applied no additional scaling (scaling_factor = "NONE") and leveraged parallel processing via BiocParallel::MulticoreParam(workers = 4) to improve efficiency. To generate test data, completely independent of training data, we prepared pseudobulk data we set ncells = 200 to select 200 cells per pseudobulk sample, and nsamples = 500 to create 500 bulk samples. We applied no additional scaling (scaling_factor = "NONE") and leveraged parallel processing via BiocParallel::MulticoreParam(workers = 4) to improve efficiency. Both pseudobulk gene expression data and cell fraction count were generated as output.

##### Other test and validation data generation

For mouse data, developmental data, and malignant data, the scRNAseq input data were divided into two parts in 50:50 cells per cell type. Using the first part for training pseudobulk generation, we began by creating a dataset using the dataset() function, providing filtered annotations, count matrix, and TPM matrix. We used the simulate_bulk() function to simulate pseudobulk data, specifying scenario = "random" to introduce random variation. Additionally, we set ncells = 100 to select 100 cells per pseudobulk sample, and nsamples = 1000 to create 1000 bulk samples. We applied no additional scaling (scaling_factor = "NONE") and leveraged parallel processing via BiocParallel::MulticoreParam(workers = 4) to improve efficiency. Both pseudobulk gene expression data and cell fraction count were generated as output. The other 50% data were used to create an independent pseudobulk dataset. To generate test data, completely independent of training data, we prepared pseudobulk datasets with ncells = 200 to select 200 cells per pseudobulk sample, and nsamples = 500 to create 500 bulk samples. We applied no additional scaling (scaling_factor = "NONE") and leveraged parallel processing via BiocParallel::MulticoreParam(workers = 4) to improve efficiency.

For test and validation on pseudobulk data generated from reannotated scRNAseq tumor datasets, we have taken the brain scRNAseq data and we began by creating a dataset using the dataset() function, providing filtered annotations, count matrix, and TPM matrix. We used the simulate_bulk() function to simulate pseudobulk data, specifying scenario = "random" to introduce random variation. Additionally, we set ncells = 900 to select 900 cells per pseudobulk sample, and nsamples = 10000 to create 10000 bulk samples. We applied no additional scaling (scaling_factor = "NONE") and leveraged parallel processing via BiocParallel::MulticoreParam(workers = 5) to improve efficiency.. Both pseudobulk gene expression data and cell fraction count were generated as output. To generate test data, completely independent of training data, we prepared pseudobulk datasets with

1. ncells = 30 to select 30 cells per pseudobulk sample, and nsamples = 1000 to create 1000 bulk samples.
2. ncells = 300 to select 300 cells per pseudobulk sample, and nsamples = 1000 to create 1000 bulk samples.
3. ncells = 900 to select 900 cells per pseudobulk sample, and nsamples = 1000 to create 1000 bulk samples.

We applied no additional scaling (scaling_factor = "NONE") and leveraged parallel processing via BiocParallel::MulticoreParam(workers = 4) to improve efficiency.

We also prepared pseudobulk to check if our model is able to capture the absence of cell types in pseudobulk samples efficiently. For that, we created two pseudobulk cohort, one with malignant, and all three CSCs blacklisted; and another with all the other cell types blacklisted. The same simulation parameters were used, with random variations in cell type proportions while excluding blacklisted cells from each simulation. To make it tough, we merged these two datasets to create a merged dataset.

##### Tumor pseudobulk data generation

To create training data suitable for tumor deconvolution, it's crucial to simulate realistic pseudobulk samples that align with the biological context of actual tumor biopsies. In real biopsies, a sample is only deemed adequate if it contains at least 10% malignant cells, reflecting a realistic tumor environment. During deconvolution, excluding malignant cells—as done in approaches like EcoTyper[108]—can lead to misleading results. For example, if a biopsy sample contains 80% malignant cells and these are removed, the majority of RNA-seq signals (80%) would become noise, overshadowing the true signals (20%) from other cell types. Moreover, purely random pseudobulk generation fails to capture the heterogeneity of cellular compositions in the tumor microenvironment[109–111]. This complexity arises from varying levels of immune infiltration, stromal cells, and cancer cells within different tumor samples. Additionally, not every tumor will contain all cell types, so a robust deconvolution model should be able to predict the absence of certain cell types in real samples. By incorporating these considerations, the simulated training data better reflects the complexities of tumor biopsies and improves the reliability of deconvolution algorithms. To generate pseudobulk data, we utilized the SimBu R package. Initially, we prepared input data, including annotation files for cell and gene information, a count matrix (readMM), and a Transcripts Per Million (TPM) matrix. Cells and genes were matched across datasets, and matrices were converted to sparse formats for efficient processing. The SimBu::dataset() function was used to create a structured dataset.

We simulated pseudobulk samples by applying SimBu::simulate_bulk() under different scenarios:

1. Weighted Scenario: We simulated varying cell-type weights (weighted_amounts) across samples by specifying a weighted cell type (e.g., "Malignant"). Multiple pseudobulk samples were generated using this scenario with a specified number of cells per sample (ncells = 900) and the number of samples per weight (nsamples_per_weighted = 3). This approach introduced gradual changes in the weighted amount to explore biological heterogeneity.

2. Blacklist-Based Simulation: We created pseudobulk samples by excluding specific cell types defined in a blacklist_list. The same simulation parameters were used, with random variations in cell type proportions while excluding blacklisted cells from each simulation.

3. Random Scenario: We additionally generated pseudobulk samples by using a random cell distribution strategy with specified whitelisted cell types. This simulation served as a control to capture unbiased cell type distributions.

After creating multiple pseudobulk datasets, we utilized the SimBu::merge_simulations() function to combine all simulated data into a comprehensive dataset. We extracted pseudobulk count matrices and cell fractions from the merged dataset using the SummarizedExperiment package and saved the results as CSV files for downstream analysis.

#### Deconvolution model optimization and comparison

##### Attention mechanism

Attention mechanisms are neural network components that allow models to selectively focus on different parts of the input data by assigning varying importance (weights) to specific elements[112]. This selective focusing improves the model's ability to identify and extract critical information. Attention is especially beneficial in complex tasks where specific features or patterns in the data play a larger role in accurate predictions. In the context of tissue deconvolution, where the input comprises pseudobulk gene expression data—a mixture of gene expression profiles from multiple cell types—the attention mechanism is crucial for distinguishing and emphasizing signals specific to different cell types. It enables the model to assign higher importance to genes that are key contributors to the expression profile of each cell type, which enhances the separation of mixed signals. This selective weighting aligns with biological principles, where certain genes are more relevant for distinguishing cell types within a heterogeneous tissue sample.

We experimented with the integration of the attention mechanism at different locations within our model: (i) at the encoder's initial layer, (ii) after the first layer of the encoder, and (iii) within the latent space. Both (ii) and (iii) configurations performed similarly, indicating that the attention mechanism effectively extracted critical features regardless of its position. The model’s ability to maintain accuracy with both configurations, even as the number of neurons decreased in subsequent layers, suggests that the attention mechanism successfully captured key patterns in the gene expression data at both stages. We chose to keep the attention mechanism after the first encoder layer. This placement allows the attention mechanism to work with refined features generated by the initial layer, effectively balancing model complexity and interpretability.

We also experimented with different numbers of attention heads (n), testing configurations with n = 1, 2, 4, 6, and 8. Our results showed that using 8 attention heads yielded the best accuracy, as it enabled the model to capture more complex interactions and dependencies within the gene expression data. Multi-head attention allows the model to look at the input from multiple perspectives, enhancing its ability to differentiate between cell types.

The suboptimal performance of the attention mechanism at the initial layer of the encoder can be attributed to the lack of sufficient feature extraction at that stage. Placing attention after the first layer, which already transforms the input data into a more informative intermediate representation, allows the model to apply attention more effectively to refined features. This intermediate representation acts as a pre-filtered and encoded version of the input, enabling the attention mechanism to better distinguish between informative and non-informative features. The comparable performance of the attention mechanism in the latent space and after the first encoder layer suggests that the refined representation at either stage contains enough information for attention to effectively differentiate cell type-specific signals. Placing the attention layer after the first encoder layer allows it to operate on a more structured and informative representation of the input data, ensuring efficiency without sacrificing interpretability. Additionally, keeping the attention mechanism after the first layer simplifies the model architecture while preserving high accuracy.

##### Gradient descent and GEP adaptive prediction

The generation of the Gene Expression Profile (GEP) involves training a deep learning model to predict how different types of cells contribute to the observed gene expression levels in a sample. Let’s imagine we have data from a mixture of several types of cells, and we want to find out which genes are most active in each cell type. To do this, we start with an initial guess of how genes behave in different cell types and gradually improve this guess using an algorithm called gradient descent. Let’s think of gradient descent like finding our way down a hill in the dark. We take small steps in the direction that makes the most progress (downhill) until we reach the lowest point (best fit). In our case, the "lowest point" is when our model accurately predicts the observed gene expression data using the estimated contributions from each cell type.

In our model, we aim to estimate the Gene Expression Profile (GEP) matrix, denoted as G, which represents the contribution of each gene across different cell types. This matrix G is initialized randomly and is iteratively updated to minimize a loss function based on the observed and predicted pseudobulk data. At the start of model training, the GEP matrix is initialized using the following command within the model definition:

```
self.gep_matrix = nn.Parameter(torch.randn(input_dim, output_dim))
```

Here, torch.randn(input_dim, output_dim) generates a matrix of random values with dimensions corresponding to the number of genes (input_dim) and the number of cell types (output_dim). This random initialization serves as our initial guess for the GEP matrix, G, which will be refined over successive iterations.

By defining self.gep_matrix as a nn.Parameter, we signal to the PyTorch framework that this matrix is a trainable component of our model. This approach leverages automatic differentiation provided by PyTorch's autograd module. Essentially, this computes the partial derivatives of the total loss with respect to each parameter using the chain rule., enabling the framework to compute gradients and update the gep_matrix during training. The initialization of gep_matrix using random values follows standard practice in deep learning, where weights and parameters are typically initialized randomly to break symmetry and avoid biases in learning. This allows the model to explore diverse paths during optimization, ultimately enhancing training efficiency and convergence. The main role of the GEP matrix in this model is to capture cell-type-specific gene expression patterns. Allowing these patterns to be learned during training from scratch (i.e., starting with random values) means that the model has the flexibility to adapt these patterns based on the specific characteristics of the dataset.

In our implementation, we utilize the Adam optimizer to perform this gradient descent step. Adam is a robust optimization algorithm that incorporates the advantages of both the Adaptive Gradient Algorithm (AdaGrad) and Root Mean Square Propagation (RMSProp). It adapts the learning rate for each parameter individually, thereby improving convergence speed and model performance.

The gradient descent algorithm aims to minimize $\mathcal{L}_{recon}$ by iteratively updating the values in the GEP matrix G. At each step t, the values of G are adjusted according to the gradient of the loss function with respect to G:

$$G_{t+1} = G_t - \eta . \frac{\partial \mathcal{L}}{\partial G_t}$$

The core objective of our method is to reconstruct the observed pseudobulk data using the estimated cell fractions and the gene expression profile (GEP) matrix. Mathematically, this reconstruction can be formulated as:

$$\hat{X} = C \, . \, G^T$$

Where:

- C represents the matrix of predicted cell fractions for each sample.
- $G^T$ denotes the transpose of the GEP matrix indicating the gene contributions for each cell type.
- $\hat{X}$ denotes the reconstructed pseudobulk.

After several iterations of this process, the GEP matrix converges to a set of values that best represents the contributions of each gene in each cell type. The resulting matrix effectively captures the gene expression signatures specific to each cell type, providing meaningful insights into the underlying biology of the cell mixture.

##### Latent space optimization

To optimize the dimensionality of the latent space in our model, we conducted systematic testing with various dimensional configurations: 64, 128, 256, 512, and 1024 dimensions. The choice of these specific dimensions was driven by a balance between model complexity and expressiveness, ensuring the latent space is sufficiently rich to capture cell-type-specific gene expression signatures without introducing unnecessary overfitting or computational overhead.

In machine learning models, especially those involving matrix factorization or representation learning tasks, the dimensionality of the latent space plays a crucial role in capturing relevant patterns. A lower-dimensional latent space, such as 64 or 128, may impose constraints on the model, forcing it to learn more compact and generalized representations. On the other hand, larger dimensions, 1024, can offer increased flexibility to capture subtle variations in gene expression but may risk capturing noise or leading to overfitting if the increase in parameters isn't justified by the data.

To determine the optimal latent space dimension, we evaluated the model’s performance using a validation set, measuring metrics such as the reconstruction error of the pseudobulk data, CCC values, and generalization on unseen datasets. Each dimensional configuration was assessed to understand the trade-off between model complexity and predictive performance, with attention paid to both accuracy and computational efficiency. This rigorous evaluation helped us find the optimal dimension that balances model expressiveness and stability, thereby improving the accuracy of our Gene Expression Profile (GEP) matrix reconstruction.

##### Hyperparameter and loss optimization

In optimizing the hyperparameters and losses for our model, we explored a comprehensive set of parameters crucial for balancing model training, regularization, and performance on biological data. For the learning rate (lr), a range of [0.1, 0.01, 0.001, 0.0001, 0.00001] was chosen to cover various magnitudes of updates to model weights. A high initial learning rate can speed up convergence, while smaller rates can help fine-tune the model and avoid oscillations near minima.

We incorporated varying dropout rates between [0.1, 0.2, 0.3, 0.4, 0.5] to regularize the model and prevent overfitting, particularly when dealing with potentially sparse or noisy biological data. The dropout mechanism randomly deactivates a proportion of neurons during training, forcing the model to generalize beyond individual neurons' dependencies. This is critical in scenarios where our deconvolution model must generalize across diverse datasets with varying signal-to-noise ratios.

The key loss components include five specific coefficients (l1 to l5), each contributing to different objectives within the overall loss function. For each coefficient, we tested a range of values: [1, 0.5, 0.1, 0.05, 0.01]. The parameter l1 governs the weight of the KL divergence loss[113], which aims to maintain similarity between predicted and prior cell type distributions, thereby grounding our predictions in biological priors. The coefficient l2 controls the loss associated with reconstructing the pseudobulk RNA-seq signal based on the learned signature matrix, representing the biological relevance of captured cell-type-specific profiles. The coefficient l3 manages the reconstruction loss between pseudobulk RNA-seq profiles and predicted gene expression profiles (GEPs), maintaining the coherence of predictions with observed data. Finally, l4 controls the alignment loss between the predicted GEP and the signature matrix, ensuring consistency between the signature matrix and the GEP representation.

To identify optimal hyperparameter settings, a grid search method was employed, systematically exploring all combinations of hyperparameters and evaluating each on a validation set. This approach allowed us to assess how each combination influences the model’s ability to reconstruct cell fractions while maintaining biological relevance. The final selected parameters balanced the multiple objectives of signal reconstruction, noise robustness, and biological fidelity, enabling the model to accurately capture cellular heterogeneity within tumor microenvironments. This rigorous optimization approach ensures the model's robustness in predicting complex cellular compositions in varying tumor biopsy conditions.

##### Comparison with other deconvolution methods

For deconvolution performance on pseudo-bulk and real bulk datasets with known ground truth, we benchmarked TAPE[114], Scaden[115], RNAsieve[116], CIBERSORTx[117], DWLS[118], Bisque[119], and EPIC[120]. The details of the benchmarking procedures are explained below, and the hyperparameter tuning information can be found in Supplementary Informations.

1. For TAPE, we evaluated its performance on a PyTorch-based version that we implemented. The training hyperparameters for the PyTorch-based TAPE were set according to the original publication and source code.
2. For Scaden, we evaluated its performance on a PyTorch-based version that we implemented. The training hyperparameters for the PyTorch-based Scaden were set according to the original publication and source code. Despite our efforts to replicate the original version closely, there were some differences, such as the loss plot and deconvolution performance on the SDY67 dataset, likely due to the different deep learning backends (Keras vs. PyTorch). However, as the

discrepancies were minor, the PyTorch implementation of Scaden was deemed acceptable. The results presented in the main section are primarily based on the Keras version, with only the performance comparison using different conducted with the PyTorch version.

3. For RNA-Sieve, due to the lack of detailed documentation, we ran it according to its example code. RNA-Sieve infers cell type proportions from bulk RNA-seq data using a composite likelihood model based on gene expression moments derived from scRNA-seq. The method leverages the central limit theorem to approximate the gene expression distribution in bulk samples as a mixture of normal distributions, accounting for the means, variances, and cell type proportions. A custom maximum likelihood estimation procedure, combined with alternating optimization and gene filtering, ensures robustness across different experimental platforms. Joint deconvolution across multiple samples further improves statistical accuracy. The documentation provided by RNA-Sieve was not sufficient for reproducibility, so we had to re-implement it on our own. We first validated its performance on pseudo-bulk data using the authors' original dataset. While it performed well on simulated data, it could not replicate the reported results on Newman’s and Monaco’s datasets.
4. For CIBERSORTx (CSx), We have rewritten the algorithm in Python from scratch, ensuring its performance remained consistent with the original implementation. CIBERSORT deconvolutes bulk RNA-seq data by leveraging a support vector regression (SVR) model trained on gene expression signatures of distinct cell types. This allows the estimation of cell type proportions in mixed samples by minimizing the difference between the observed bulk expression and a linear combination of the reference profiles. The method includes a gene filtering step based on variability and incorporates bootstrapping to provide robust estimates and confidence intervals for the inferred proportions. Despite reimplementing it in Python, the performance and accuracy were unaffected, yielding reliable deconvolution results. During comparison, we first generated a signature matrix using a single-cell profile, then deconvolved the corresponding bulk data with batch correction, keeping other settings as default.
5. For DWLS, we utilized the core R functions and packages to create signature matrices and deconvolve the target pseudo-bulk and real bulk datasets. To ensure the correctness of our implementation, we adhered to the intestine stem cell example from the DWLS manual. Since DWLS deconvolves samples individually, we used a for-loop to handle larger samples. To optimize DWLS performance, we used the Seurat flavor for the pseudo-bulk test and the MAST flavor for the real bulk test. Additionally, to improve stability across all bulk samples, we employed support vector regression (nu-SVR) for initial estimation instead of ordinary least square regression.
6. For Bisque, we installed the R package BisqueRNA and followed the author’s example with default settings. In its "Reference-based decomposition" mode,

Bisque automatically filters low-variance genes and uses the remaining genes for decomposition. We provided all the genes without specifying marker genes.

7. For EPIC, we used its web interface with our own reference gene expression data for each case.

Both CIBERSORT and TAPE models are recognized for their ability to generate gene expression profiles (GEPs) alongside predicting cell fractions. This GEP output is crucial for enhancing the interpretability of these models. To evaluate this aspect, we conducted a comparative analysis of our GEPs against those generated by CIBERSORT and TAPE. The comparison focused on two key areas. First, we assessed how well each set of GEPs could reconstruct the original pseudobulk expression when provided with known cell fractions. The accuracy of this reconstruction was quantified using the Concordance Correlation Coefficient (CCC) between the original and reconstructed pseudobulk profiles. Additionally, we compared the GEPs by examining their alignment with experimentally validated cell type-specific markers, providing further insight into the biological relevance and specificity of each model's GEP output.

Malignant cells lack well-defined specific gene expression markers, making it challenging to directly validate their gene expression profiles (GEPs). However, telomerase expression is often elevated in cancerous cells and can serve as a potential marker to evaluate the biological relevance of malignant cell GEPs. In our comparison of GEPs, alongside evaluating the reconstruction of the original pseudobulk using known cell fractions and comparing with experimentally validated cell-type specific markers, we also examined the expression of telomerase as an indicator of malignant cell validity. Specifically, we checked for elevated telomerase expression levels within the malignant cell GEPs. This additional analysis provides a biological context to assess the accuracy of malignant cell profiling, ensuring that the predicted GEPs not only match overall pseudobulk but also reflect expected cancer-associated gene expression characteristics.
